# Large genetic diversity and strong positive selection in F-box and GPCR genes among the wild isolates of *Caenorhabditis elegans*

**DOI:** 10.1101/2020.07.09.194670

**Authors:** Fuqiang Ma, Chun Yin Lau, Chaogu Zheng

## Abstract

The F-box and chemosensory GPCR (csGPCR) gene families are greatly expanded in nematodes, including the model organism *Caenorhabditis elegans*, compared to insects and vertebrates. However, the intraspecific evolution of these two gene families in nematodes remain unexamined. In this study, we analyzed the genomic sequences of 330 recently sequenced wild isolates of *C. elegans* using a range of population genetics approaches. We found that F-box and csGPCR genes, especially the *Srw* family csGPCRs, showed much more diversity than other gene families. Population structure analysis and phylogenetic analysis divided the wild strains into eight non-Hawaiian and three Hawaiian subpopulations. Some Hawaiian strains appeared to be more ancestral than all other strains. F-box and csGPCR genes maintained a great amount of the ancestral variants in the Hawaiian subpopulation and their divergence among the non-Hawaiian subpopulations contributed significantly to population structure. F-box genes are mostly located at the chromosomal arms and high recombination rate correlates with their large polymorphism. Moreover, using both neutrality tests and Extended Haplotype Homozygosity analysis, we identified signatures of strong positive selection in the F-box and csGPCR genes among the wild isolates, especially in the non-Hawaiian population. Accumulation of high-frequency derived alleles in these genes was found in non-Hawaiian population, leading to divergence from the ancestral genotype. In summary, we found that F-box and csGPCR genes harbour a large pool of natural variants, which may be subjected to positive selection. These variants are mostly mapped to the substrate-recognition domains of F-box proteins and the extracellular and intracellular regions of csGPCRs, possibly resulting in advantages during adaptation by affecting protein degradation and the sensing of environmental cues, respectively.

**Significance statement:** The small nematode *Caenorhabditis elegans* has emerged as an important organism in studying the genetic mechanisms of evolution. F-box and chemosensory GPCR proteins are two of the largest gene families in *C. elegans*. However, their intraspecific evolution within *C. elegans* was not studied before. In this work, using the nonsynonymous SNV (single nucleotide variant) data of 330 *C. elegans* wild isolates, we found that F-box and chemosensory GPCR genes showed larger polymorphisms and stronger positive selection than other genes. The large diversity is likely the result of rapid gene family expansion, high recombination rate, and gene flow. Analysis of subpopulation suggests that positive selection of these genes occurred most strongly in the non-Hawaiian population, which underwent a selective sweep possibly linked to human activities.

## Introduction

*C. elegans* genome contains over 350 F-box genes, compared to ∼69 in human genome (Kipreos and Pagano 2000; Thomas 2006). This great expansion of the F-box gene family is the result of tandem gene duplication, which has also been observed in plants (Xu, et al. 2009). F-box genes code for proteins sharing the F-box domain, a 42-48 amino acid-long motif that binds to Skp1 (S-phase kinase-associated protein 1) proteins during the assembly of the SCF (Skp1-Cullin-F-box) E3 ubiquitin ligase complexes, which ubiquitinate protein substrates and target them for degradation. F-box proteins also contain substrate-binding domains, including FOG-2 homology (FTH) domain, F-box associated (FBA) domain, Leucine-rich repeats (LRR) and WD40 repeats, which recruit the substrate proteins to the E3 ubiquitin ligase (Kipreos and Pagano 2000). F-box genes and the SCF complex-mediated protein degradation have diverse functions in *C. elegans*, including the regulation of lifespan (Ghazi, et al. 2007), developmental timing (Fielenbach, et al. 2007), sex determination (Jager, et al. 2004), and neuronal differentiation (Bounoutas, et al. 2009). The role of F-box proteins in the evolution of *Caenorhabditis* species has been noticed before in the study of sex determination and the rise of hermaphroditism. For example, through convergent evolution, *C. elegans* and *C. briggsae* independently evolved the hermaphroditic reproduction system by using two different F-box genes (*fog-2* and *she-1*, respectively) to suppress the translation of *tra-2* mRNA and promote spermatogenesis (Guo, et al. 2009). The intraspecific variation of F-box genes and their contribution to adaptation within *C. elegans* have not been studied.

Chemoreception is a major way for the nematodes to sense environmental cues and is mediated by the chemosensory-type seven-transmembrane G-protein-coupled receptors (csGPCRs). The *C. elegans* genome contains more than 1,300 csGPCR genes (Thomas and Robertson 2008), an exceptionally large number given the small size of its nervous system (302 neurons in adult hermaphrodites). The csGPCR genes can be divided into four superfamilies and families (in parentheses): *Str* (*srd, srh, sri, srj*, and *str*), *Sra* (*sra, srab, srb*, and *sre*), *Srg* (*srg, srt, sru, srv, srx*, and *srxa*), and *Solo* (*srw, srz, srbc, srsx*, and *srr*) (Vidal, et al. 2018). Evidence of extensive gene duplication and deletion and intron gain and loss were found in the *srh* genes among species in the *Caenorhabditis* genus, suggesting rapid interspecific evolution (Robertson 2000). Several of the csGPCRs were found to be essential for sensing some odors and pheromones (Sengupta, et al. 1996; Kim, et al. 2009; Park, et al. 2012), but the function of most csGPCRs is unknown. The expansion of the csGPCR gene families and their roles in environmental sensing strongly suggest their involvement in evolution, but the evidence for intraspecific positive selection is missing.

Thanks to the sampling efforts in the past, a collection of ∼330 wild isolates of *C. elegans* have been obtained and sequenced (Crombie, et al. 2019; Stevens, et al. 2019). Their genomic sequences were recently made available (Cook, et al. 2017), providing an important resource for understanding the intraspecific evolution of *C. elegans*. Here we analyzed the SNVs among the 330 wild isolates of *C. elegans* and compared the nucleotide diversity of genes belonging to different gene families. We found that the F-box and the csGPCR genes showed much larger diversity than an average gene. Population structure analysis divided the wild strains into eight non-Hawaiian and three Hawaiian subpopulations. F-box and csGPCR genes maintained a large amount of potentially ancestral variant sites in the Hawaiian strains and their divergence among the eight non-Hawaiian groups contributed significantly to population structure. Given their location at mostly the chromosomal arms, high recombination rate might have contributed to the large diversity of these genes. Furthermore, both neutrality tests and Extended Haplotype Homozygosity analysis identified signs of strong positive selection in the F-box and csGPCR genes among the wild isolates, especially in the non-Hawaiian population; derived alleles of these genes might have altered gene functions, leading to selective advantages. In summary, our systematic analysis suggests that F-box and csGPCR genes harbour a large pool of natural variants, which were subjected to positive selection; such selection may have contributed to the recent selective sweep and adaptive evolution of the wild *C. elegans* population.

## Results

### Large polymorphisms in F-box and chemosensory GPCR genes among *C. elegans* wild isolates

From the sequencing data of 330 distinct isotypes of *C. elegans* wild strains (VCF files of 20180527 release on CeNDR), we identified in total 2,493,687 SNVs, including 271,718 SNVs synonymous and 266,004 nonsynonymous SNVs (supplementary fig. S1A, Supplementary Material online). By analyzing the distribution of the variants across 20,222 protein-coding genes, we found that 1,143 genes with average CDS length of 0.6 kb (genomic average is 1.2 kb) had no nonsynonymous mutations or small indels in CDS; within the 1,143 genes, 302 with the average gene length of 1.4 kb (genomic average is 3.1 kb) did not have any SNVs or small indels in any of the CDS, intron, and UTR regions. The absence of coding variations in these genes may be explained by their small size, enrichment in regions with low recombination rate (e.g. *Rho* = 0 for 969 of the 1143 genes; see below for the genomic distribution of *Rho*), and possible purifying selection (many of them are involved in cell division and germline development; supplementary fig. S3, Supplementary Material online).

Next, we calculated nucleotide polymorphism (*Pi*) for each protein-coding gene. We found that *Pi* is significantly larger in noncoding regions (introns and UTRs) compared to those in synonymous and nonsynonymous sites in coding regions (supplementary fig. S1C, Supplementary Material online). For nonsynonymous SNVs, we found that the F-box and csGPCR genes have much larger diversity than average genes (fig. 1). For example, among the 235 genes whose *Pi* is bigger than 0.01, 46 of them are F-box genes, indicating an over ten-fold enrichment (fig. 1A). Compared to other gene families like the transcription factor (TF) genes (891) and the protein kinase genes (402), F-box genes (336) and the csGPCRs (1301) on average have significantly bigger *Pi* (fig. 1B and C; significance by a non-parametric Wilcoxon’s test). Large genetic diversity among the wild isolates hints that the F-box and csGPCR genes might contribute to the adaptation of *C. elegans* in the natural environment.

**Figure 1.**
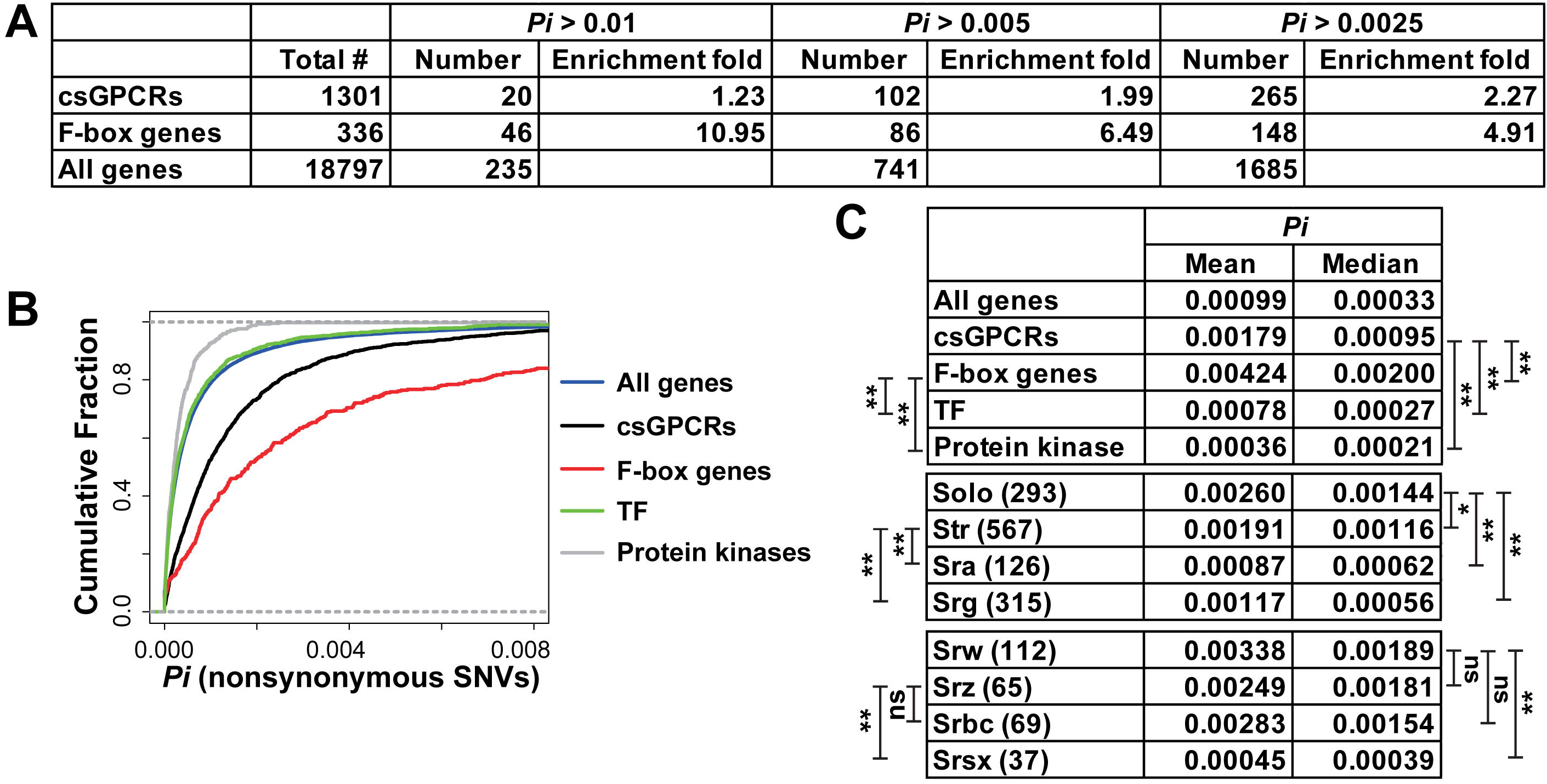
Large genetic polymorphism of csGPCR and F-box genes. (A) Genes with large *Pi* for nonsynonymous SNVs tend to be enriched in csGPCR and F-box gene families. *Pi* values of individual genes can be found in Table S4. (B) The cumulative distribution of the *Pi* values for all genes, csGPCR, F-box, transcription factor (TF) and Protein kinases genes. (C) The mean and median of *Pi* for different gene families and for different csGPCR superfamilies. The number of genes are in the parentheses. For statistical significance in a non-parametric Wilcoxon’s rank-sum test, ns means not significant, a single asterisk means *p* < 0.05, and double asterisks mean *p* < 0.01. Similar annotations apply for the rest of the Figures.

The csGPCRs can be further divided into *Str, Sra, Srg*, and *Solo* superfamilies, among which the Solo superfamily genes have the biggest *Pi* (fig. 1C). Within the *Solo* superfamily, *Srw*-type csGPCRs appeared to have the largest polymorphism on average, although the mean of *Pi* is not significantly bigger than *Srz* and *Srbc* subfamilies. The large genetic diversity is correlated with the abundance of segregating sites; F-box and the *Srw* genes both have over three times more variant sites (45.7 and 44.2, respectively) than the average of all genes (14.0). In extreme cases, *srw-57* has only 1071 nucleotide in the CDS but carries 124 nonsynomymous variants; *fbxb-53* is 1020-bp long in the CDS and has 207 segregating sites.

### Genetic divergence of F-box and csGPCR genes among *C. elegans* subpopulations

We next conducted population structure analysis on the 330 wild isolates and found 3 Hawaiian and 8 non-Hawaiian subpopulations (supplementary fig. S2 and table S1, Supplementary Material online), which generally agrees with a recent study that used 276 strains and also found 11 distinct genetic groups (Crombie, et al. 2019). Phylogenetic analysis using all nonsynymous SNVs and neighbor-joining methods (Huson and Bryant 2006) showed the evolutionary relationship among the *C. elegans* wild isolates. We rooted the tree using *C. briggsae, C. remanei* and *C. brenneri* as outgroups (fig. 2A; see Materials and Methods) to show that Hawaiian strains, especially “Hawaii_1” and “Hawaii_2” groups, are genetically closer to the sister species and contain more ancestral variations than the non-Hawaiian strains. Two “Hawaii_1” strains XZ1516 and ECA701 are highly divergent from other strains. “Hawaii_3” strains cluster more closely with non-Hawaiian strains (fig. 2A) and are more admixed with non-Hawaiian subpopulations (supplementary fig. S2, Supplementary Material online) compared to “Hawaii_”1 and “Hawaii_2” strains, likely due to gene flow (supplementary fig. S4A, Supplementary Material online).

**Figure 2.**
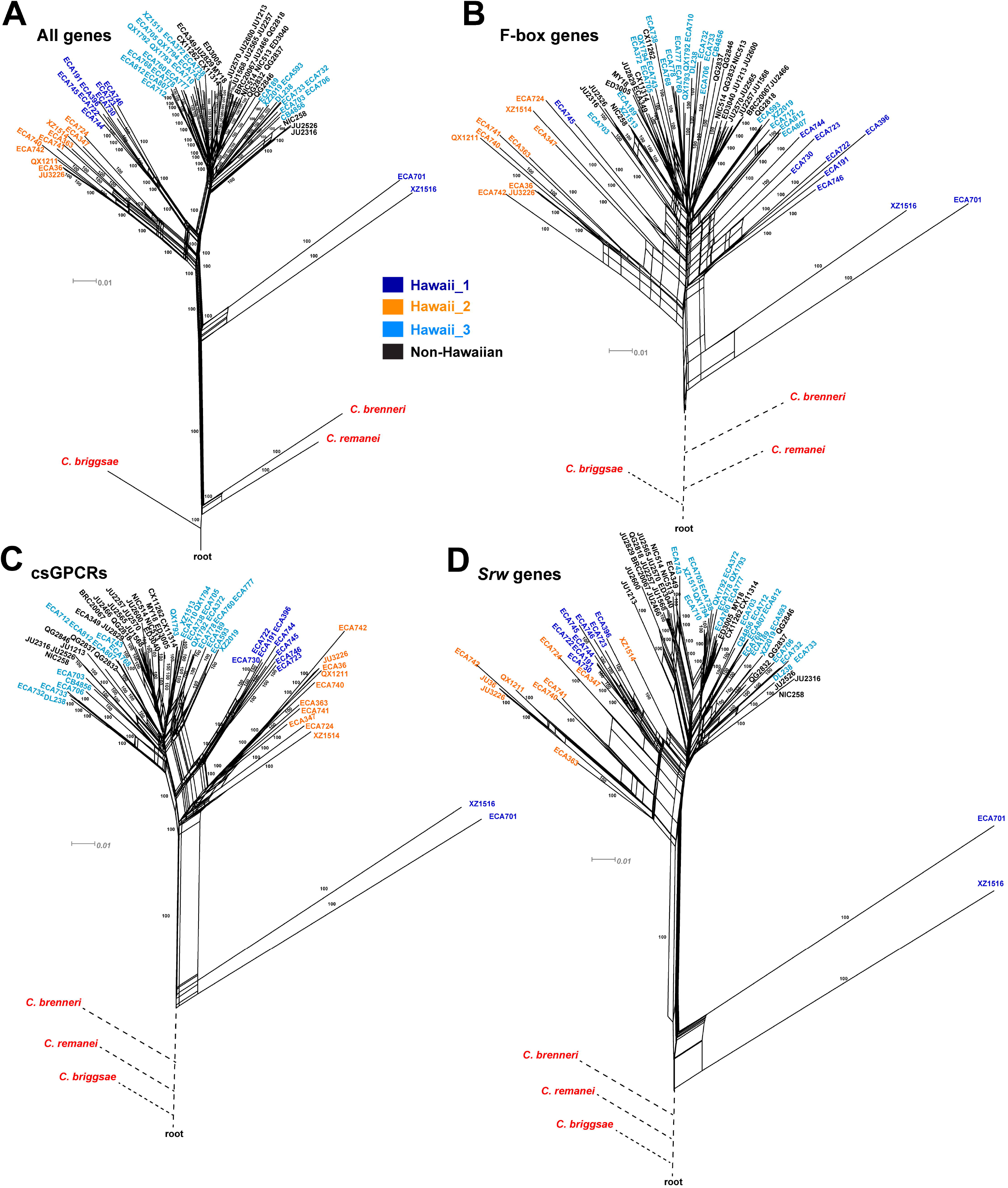
Phylogenetic relationship of the *C. elegans* wild isolates. Neighbour-joining nets plotted using the nonsynonymous SNVs of all genes (A), F-box genes (B), csGPCRs (C), or *Srw* genes (D). *C. brenneri, C. remanei*, and *C. briggsae* were used as outgroups for tree construction. Three representative non-Hawaiian strains (in black) with high ancestral population fraction were chosen from each of the eight non-Hawaiian groups. Edges are labelled with “100”, if 100% bootstrap support was attained in 1,000 bootstrap replicates. To fit the trees into one figure, some branches connecting the three outgroups and the root are manually shortened (dashed lines).

Compared to the genomic average, divergence between the Hawaiian and non-Hawaiian strains are deeper in F-box and csGPCR genes, as shown in expanded neighbor-joining net and increased phylogenetic distance (fig. 2B and C). Within csGPCRs, *Srw* genes appear to show even greater divergence among the subpopulations (fig. 2D). Moreover, the phylogenetic trees constructed using nonsynomymous SNVs of csGPCR or F-box genes had different topologies from the tree of all genes (fig. 2D). For example, looser clustering patterns and more admixture between Hawaiian and non-Hawaiian strains were observed for the F-box and *Srw* genes, suggesting that these genes may have a distinct evolutionary history than other genes.

Based on the population structure and genetic grouping, we divided the 330 wild isolates into Hawaiian (45 strains) and non-Hawaiian (285) populations (supplementary table S2, Supplementary Material online) and calculated polymorphism for the two populations using nonsynomymous SNVs. Hawaiian population showed over two-fold larger *Pi* than non-Hawaiian population across all genes (fig. 3A), which is consistent with the hypothesis that recent selective sweep reduced variation in non-Hawaiian population, while Hawaiian strains kept part of the ancenstral diversity (Cook, et al. 2017; Crombie, et al. 2019). “Hawaii_3” has lower diversity than the other two Hawaiian subpopulations, likely because “Hawaii_3” strains are genetically more similar to the non-Hawaiian strains. Interestingly, the diversity of F-box and csGPCR genes is bigger than the TF, protein kinase genes, or an average gene in both non-Hawaiian and Hawaiian populations (fig. 3A).

**Figure 3.**
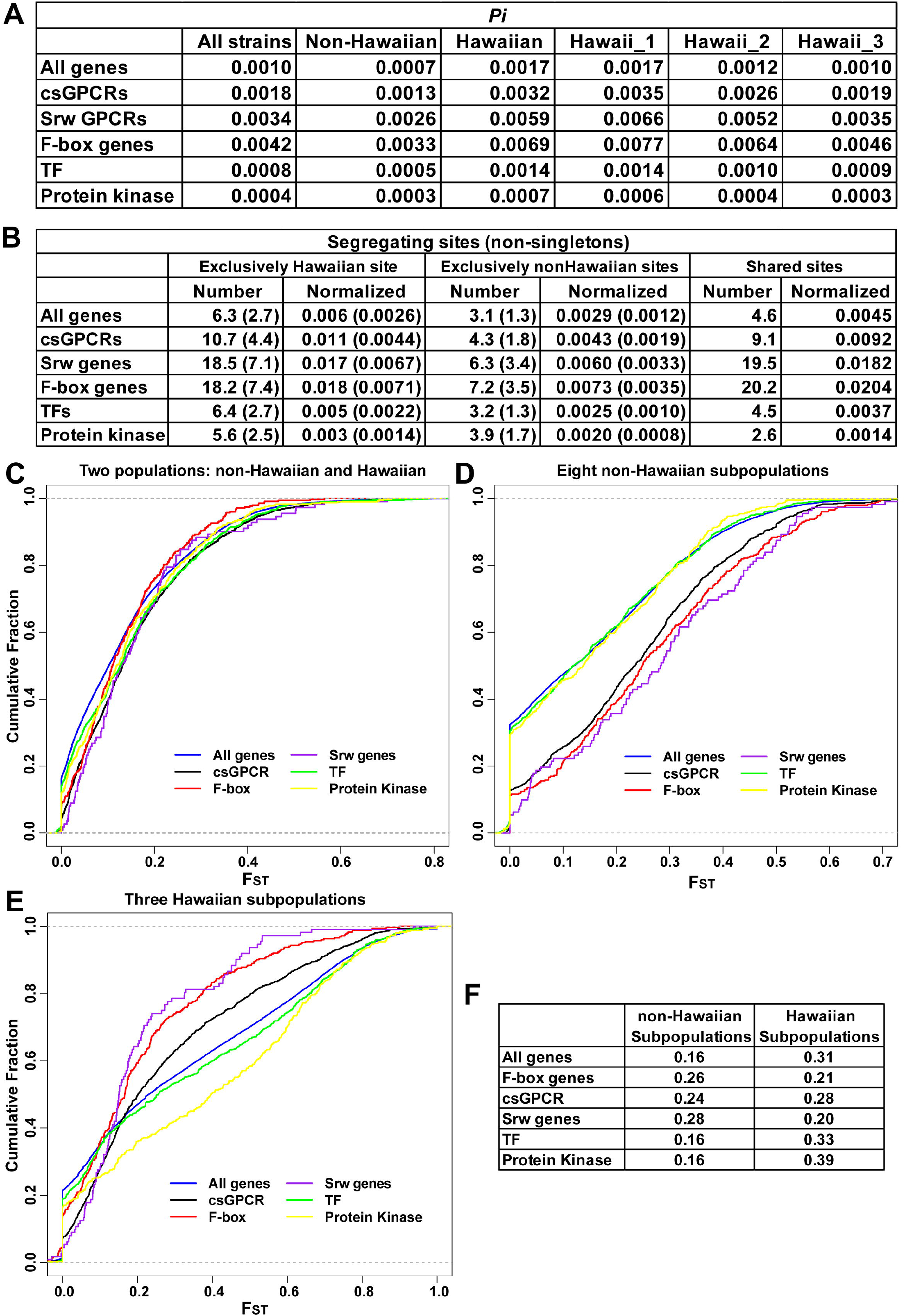
csGPCR and F-box genes contribute to the large divergence of Hawaiian strains and the differentiation among non-Hawaiian subpopulations. (A) The mean of CDS length-normalized *Pi* of all genes, csGPCRs, *Srw* genes, F-box genes, TF, and Protein kinase for non-Hawaiian and Hawaiian populations, as well as the three Hawaiian subpopulations (see the grouping in Methods). (B) The average number of segregating sites that belong to only Hawaiian or non-Hawaiian strains and the sites that are shared by Hawaiian and non-Hawaiian strains for the six gene families. The number is also normalized to the CDS length of individual genes. The number of non-singleton segregating sites are in the parentheses. (C-E) The cumulative distribution of Hudson’s F_ST_ values for different gene families between the non-Hawaiian and Hawaiian populations (C), among the eight non-Hawaiian subpopulations (D), and among the three Hawaiian subpopulations (E). (F) The average F_ST_ value of different gene families among non-Hawaiian and among Hawaiian subpopulations.

We also found that a large number (6.3 per gene on average) of segregating sites only existed in Hawaiian strains and much fewer (3.1 per gene) sites are exclusively non-Hawaiian; a significant number (4.6 per gene) of sites are shared between some Hawaiian and non-Hawaiian strains (fig. 3B). As expected, F-box and csGPCR genes have a lot more exclusively Hawaiian sites than the TF or protein kinase genes. However, they do not carry many exclusively non-Hawaiian sites, and the large diversity of the F-box and csGPCR genes in non-Hawaiian strains mostly result from the large number of sites originated from the Hawaiian population (fig. 3B). This finding supports that the Hawaiian *C. elegans* (especially the “Hawaii_1” and “Hawaii_2” groups) maintains a relatively large pool of ancestral variation, and polymorphisms in the F-box and csGPCR genes contribute significantly to this ancestral diversity. Although selective sweep removed many ancestral alleles in non-Hawaiian population, the F-box and csGPCR genes still kept a significant number of variant sites, which might be related to adaptation.

Fixation index F_ST_ is a measure for genetic difference between populations. F_ST_ for F-box and csGPCR genes were similar to other gene families when just considering Hawaiian and non-Hawaiian as two populations (fig. 3C). ∼80% of all genes have F_ST_ < 0.2. We reasoned that this may be caused by large divergence among the subpopulations within each population. When calculating for the eight non-Hawaiian subpopulations (supplementary table S3, Supplementary Material online), only ∼60% of the genes have F_ST_ < 0.2 and that F-box and csGPCR genes, especially *Srw* genes, have much higher mean F_ST_ than TF and protein kinase genes or an average gene (fig. 3D and 3F). This finding suggests that the polymorphism of F-box and csGPCR genes contribute significantly to the population structure of the non-Hawaiian strains. Their divergence among subpopulations and fixation within subpopulation may be linked to local adaptation. For example, csGPCR *srw-66* (F_ST_ = 0.76) contains 24 variants that were found in >75% of the strains in the “North_America” group and >55% of the “Europe_2” strains but not in any other non-Hawaiian groups. Similarly, F-box gene *fbxa-181* (F_ST_ = 0.69) has 14 SNVs that are found in 73% of the “Europe_6” strains and not in any other groups.

Among the three Hawaiian subpopulations, the mean F_ST_ values of F-box and csGPCR *Srw* genes appear to be significantly lower than other gene families or the genomic average (fig. 3E and 3F), which may be explained by their large diversities even within the same Hawaiian group (fig. 3A). Thus, the big variation of F-box and *Srw* genes do not seem to follow the population structure among the three Hawaiian subpopulations and they are not likely fixed within the Hawaiian groups.

Gene flow also helped shape the diversity of the F-box and csGPCR genes. F-box genes have extensive gene flow between Hawaiian and non-Hawaiian populations in both directions (supplementary fig. S4B, Supplementary Material online), which is consistent with the great number of shared segregating sites in F-box genes between the two populations (fig. 3B). On the other hand, csGPCR genes had only gene flow within the non-Hawaiian subpopulations. Interestingly, when constructing the maximum-likehood population tree for gene flow analysis, we found that the tree structure changed after removing the variants in F-box or csGPCR genes. Instead of staying as a branch outside of the eight non-Hawaiian subpopulations, the “Hawaii_3” group moved into the non-Hawaiian groups and was placed next to “Europe_2” and “North_America” (supplementary fig. S4C, Supplementary Material online). This finding supports that variations in the F-box and csGPCR genes played critical roles in distinguishing “Hawaii_3” strains from the non-Hawaiian populations and contributed significantly to intraspecific diversity.

### High recombination rate may contribute to the polymorphism of the F-box genes

We next asked whether chromosomal locations of the F-box and csGPCR genes had effects on their diversity. Most of the F-box genes are located on the arms of chromosome II (33%), III (22%), and V (26%) (fig. 4A). In contrast, protein kinase and TF genes are more evenly spread out across chromosomes. Distal regions of the chromosomes tend to have higher frequencies of recombination and larger polymorphisms than the center of chromosomes (Begun and Aquadro 1992; McGaugh, et al. 2012). Indeed, when using the entire set of 2,493,687 SNVs and the FastEPRR software (Gao, et al. 2016) to estimate the recombination rates, we observed both higher recombination rate (*Rho*) and higher variant density in the chromosomal arm regions (two distal quarters) compared to the centers (the middle half) of chromosome II, III, and V (fig. 4B). Polymorphisms for both nonsynonymous and synonymous SNVs appeared to be higher in high recombination region (*Rho* > 0) than in low recombination region (*Rho* = 0) for all genes (fig. 4C); and a positive correlation between the recombination rate and *Pi* values was observed (fig. 4D). F-box genes showed almost two-fold enrichment in high recombination region and F-box genes in regions with higher recombination rates had higher levels of polymorphism (fig. 4C). Thus, the genomic location of F-box genes in the chromosomal arms may contribute to their large genetic diversity.

**Figure 4.**
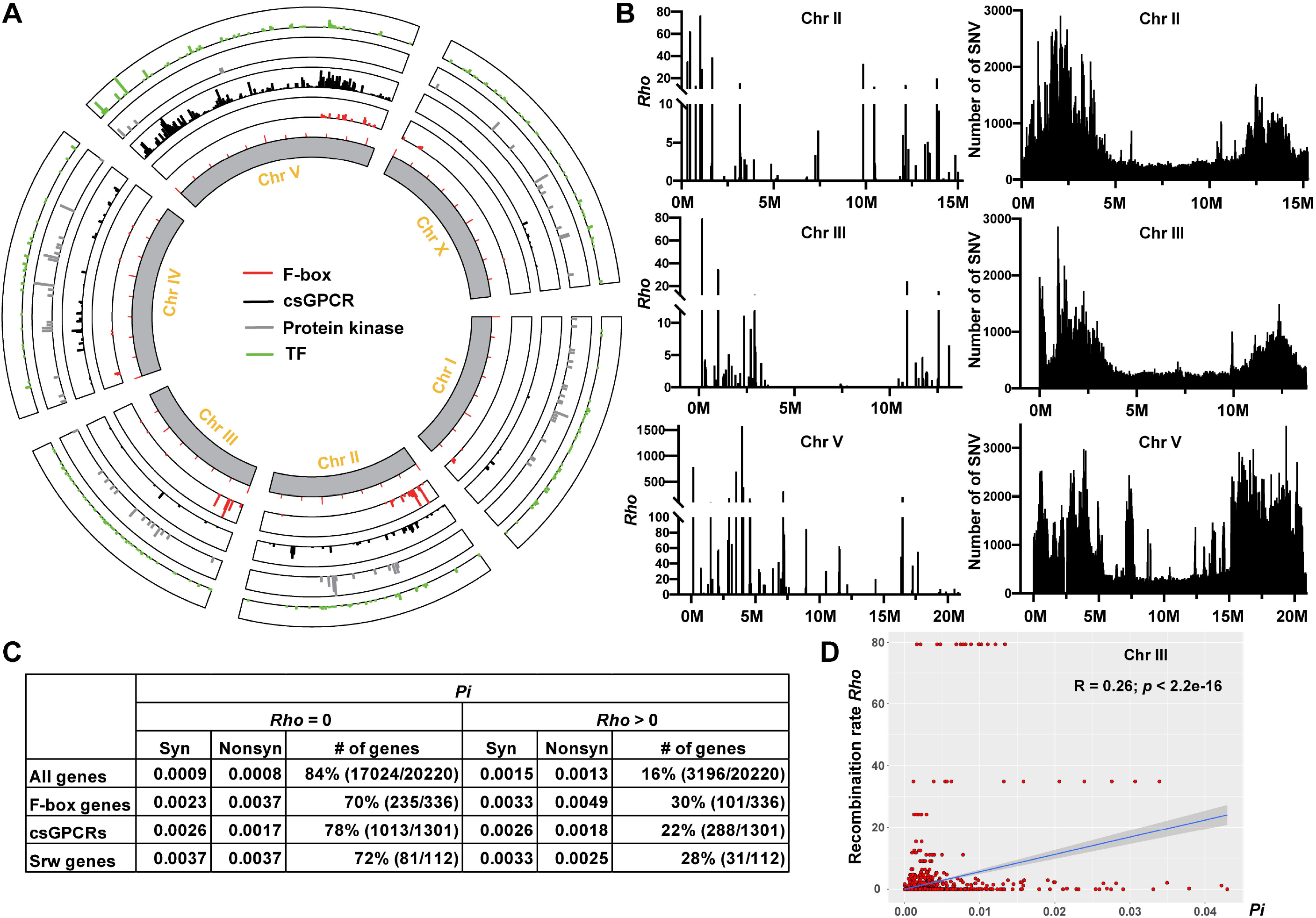
High recombination rate may contribute to the large diversity of F-box genes. (A) Genomic location of F-box, csGPCR, Protein kinase, and TF genes plotted using TBtools. (B) Recombination rates (*Rho*) and the density of SNVs across Chr II, III and V in 50-kb windows. (C) The polymorphism for synonymous and nonsynonymous SNVs in the low (*Rho* = 0) and high (*Rho* > 0) recombination regions. (D) The Pearson correlation between recombination rate and the *Pi* of all SNVs for individual genes on Chr III.

Most of the csGPCR genes are located on chromosome II (13%), IV (9%), and V (70%) (fig. 4A). Chromosome V is the biggest among the six chromosomes, has the highest variant density, and contains regions with very high recombination rates (fig. 4B). Compared to F-box genes, csGPCRs are less concentrated on the chromosomal arms. Although csGPCRs, e.g. *Srw* genes, showed enrichment in high recombination region, we did not observe increased polymorphism for csGPCRs in the high recombination region than in the low recombination region (fig. 4C).

Besides recombination, the abundance of polymorphic sites may also be the consequence of gene duplication. Chromosomal clustering of F-box and csGPCR genes indicates rapid gene family expansion through often tandem or inverted duplications (Robertson and Thomas 2006), which creates genetic redundancy and allows the accumulation of variants. Our analysis of copy number variants (CNVs) among the wild isolates supported this idea. Among the 8740 CNVs found in 5586 genes, 185 (1.99 fold enrichment) F-box genes and 552 (1.54 fold enrichment) csGPCRs carried CNVs (supplementary fig. S5A, Supplementary Material online). Moreover, the average number of CNVs per gene is also higher for F-box and csGPCR genes compared to genomic average. Thus, large genetic polymorphisms for these genes were reflected in both the abundance of SNVs and CNVs.

### Signs of strong positive selection on F-box and csGPCR genes

Previous studies hypothesized that positive selection of alleles that confer fitness advantages under human influence reduced genetic variations in *C. elegans* (Andersen, et al. 2012), but the genes under selection are unknown. Using the nonsynonymous SNVs, we performed neutrality tests and calculated Tajima’s *D* and Fay and Wu’s *H* values for every gene. The *D* value reflects the difference between expected and observed diversity (Tajima 1989) and the *H* value measures the abundance of high-frequency derived allele (Fay and Wu 2000). Negative *D* and *H* values are both indicators of selective sweep and positive selection. To calculate the *H* value, we used XZ1516 or ECA701 as the outgroup, because these two strains likely carry the most ancestral genotypes (fig. 2). *H* values calculated using the two strains as the outgroup were similar.

In the neutrality tests, we found that Tajima’ *D* were negative for the nonsynonymous SNVs for most (>85%) genes and Fay and Wu’s *H* were negative for ∼50% genes (supplementary table S4, Supplementary Material online). This finding is consistent with the chromosome-wide sweep that occurred across the genome (Andersen, et al. 2012). Interestingly, F-box and csGPCR genes are overrepresented among the genes with significantly negative *D* and *H* values (fig. 5A). For example, among the 1038 genes with *H* < −20, 260 of them are csGPCRs (3.62-fold enriched) and 67 are F-box genes (3.61-fold enriched). Gene ontology analysis consistently showed strong enrichment (> 5 fold) in genes involved in sensory perception of smell and chemical stimulus (supplementary fig. S6, Supplementary Material online).

**Figure 5.**
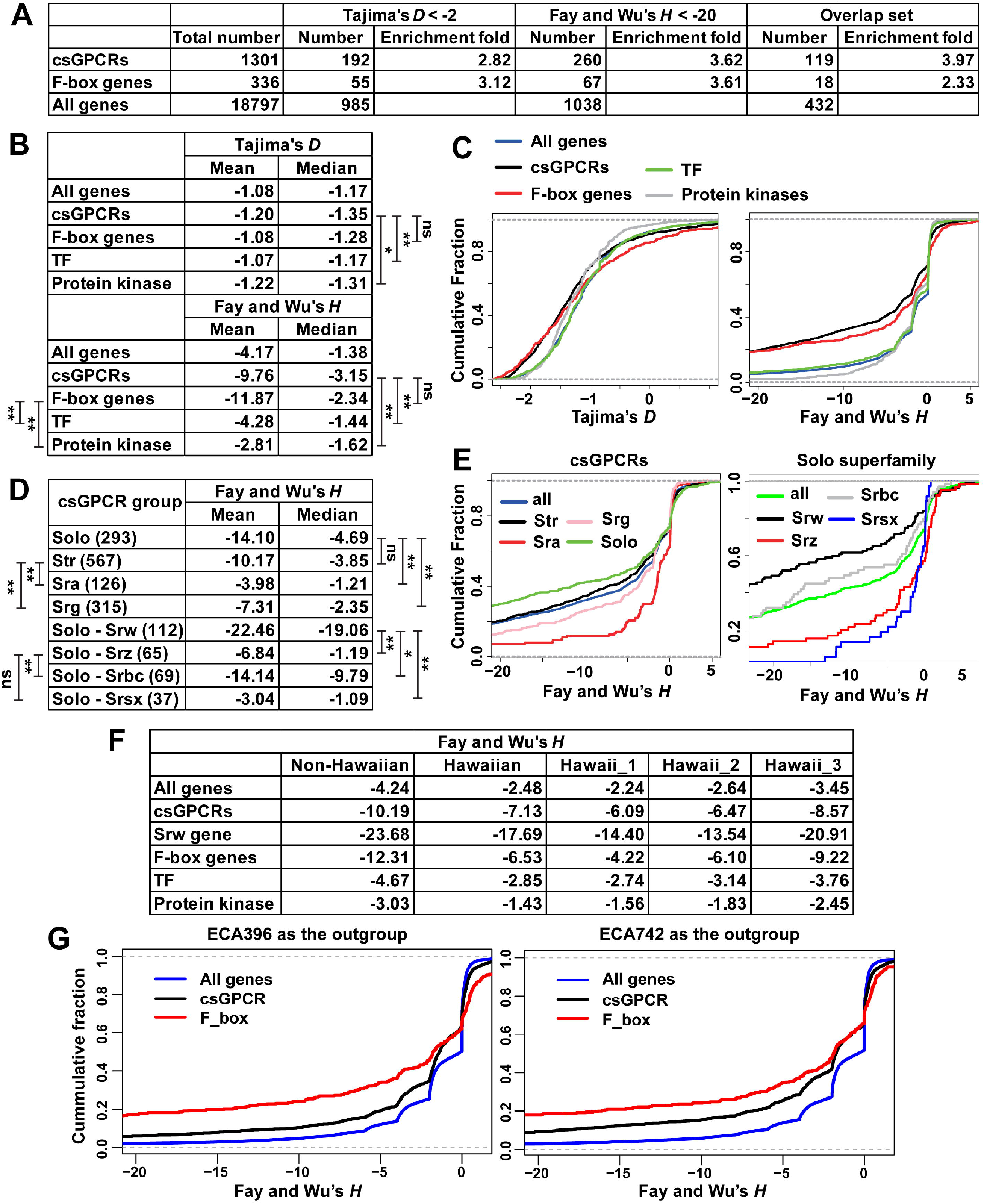
Positive selection on F-box and csGPCR gene. (A) Enrichment of csGPCR and F-box genes among the genes with Tajima’s *D* < −2 and Fay and Wu’s *H* < −20, respectively. Overlap set include genes that fits both criteria. (B) The mean and median of Tajima’s *D* and Fay and Wu’s *H* values of all genes, csGPCRs, F-box, TF, and Protein kinase. (C) The cumulative distribution of different gene families. (D) The mean and median of Fay and Wu’s *H* values of genes in csGPCR superfamilies and *Solo* gene families. The number of genes are in parentheses. (E) The cumulative distribution of genes in csGPCR subfamilies and *Solo* families. The statistical significance was determined by Wilcoxon rank-sum test. (F) The average Fay and Wu’s *H* values of all genes, csGPCRs, *Srw* genes, F-box genes, TFs, and Protein kinase for the non-Hawaiian and Hawaiian populations, as well as the three Hawaiian subpopulations. The above *H* values were all calculated using XZ1516 as the outgroup. (G) The cumulative distribution of the *H* values of all genes, csGPCR, or F-box genes calculated using ECA396 or ECA742 as the outgroup.

Compared with the TF and protein kinases genes or the genomic average, F-box and csGPCR genes have significantly lower *D* and *H* values (fig. 5B and C), suggesting that the csGPCR and F-box genes appear to be under stronger positive selection than other genes. Within the csGPCRs, Solo superfamily genes have the lowest *H* values and within the *Solo* superfamily, *Srw*-type csGPCRs have the lowest *H*, indicating that *Srw* genes may be under the strongest positive selection among all csGPCRs (fig. 5D and E). Putative functions of the *Srw* genes in sensing environmental peptides suggest they may be involved in adaptation.

Within the F-box genes, we did not observe significant difference in either *D* or *H* values or polymorphisms among the genes in *fbxa, fbxb*, and *fbxc* subfamilies (supplementary fig. S7A, Supplementary Material online). F-box proteins share an F-box domain, which complexes with Skp and Cullin to form the SCF complex that mediates protein ubiquitination and degradation. Five out of the 20 Skp-related genes in *C. elegans* (*skr-3, 4, 5, 10*, and *15*) and three out of the 6 Cullin genes (*cul-1, 3*, and *6*) have highly negative *H*, suggesting strong selective sweep (supplementary fig. S7B, Supplementary Material online). Thus, components of the ubiquitination-proteasome system (UPS) may have been co-evolving in the *C. elegans* wild isolates; genetic variations in UPS genes may alter the homeostasis of target proteins, leading to certain advantages during selection.

Another line of evidence for positive selection is the excess of nonsynonymous SNVs compared to synonymous SNVs, which is particularly obvious for F-box genes. *Pi* was bigger and *D* and *H* values were more highly negative for nonsynonymous SNVs compared to synonymous SNVs (fig. 6A); the pN/pS ratio for F-box genes is also much higher than the genomic average (fig. 6B). These results support that F-box genes are under positive selection.

**Figure 6.**
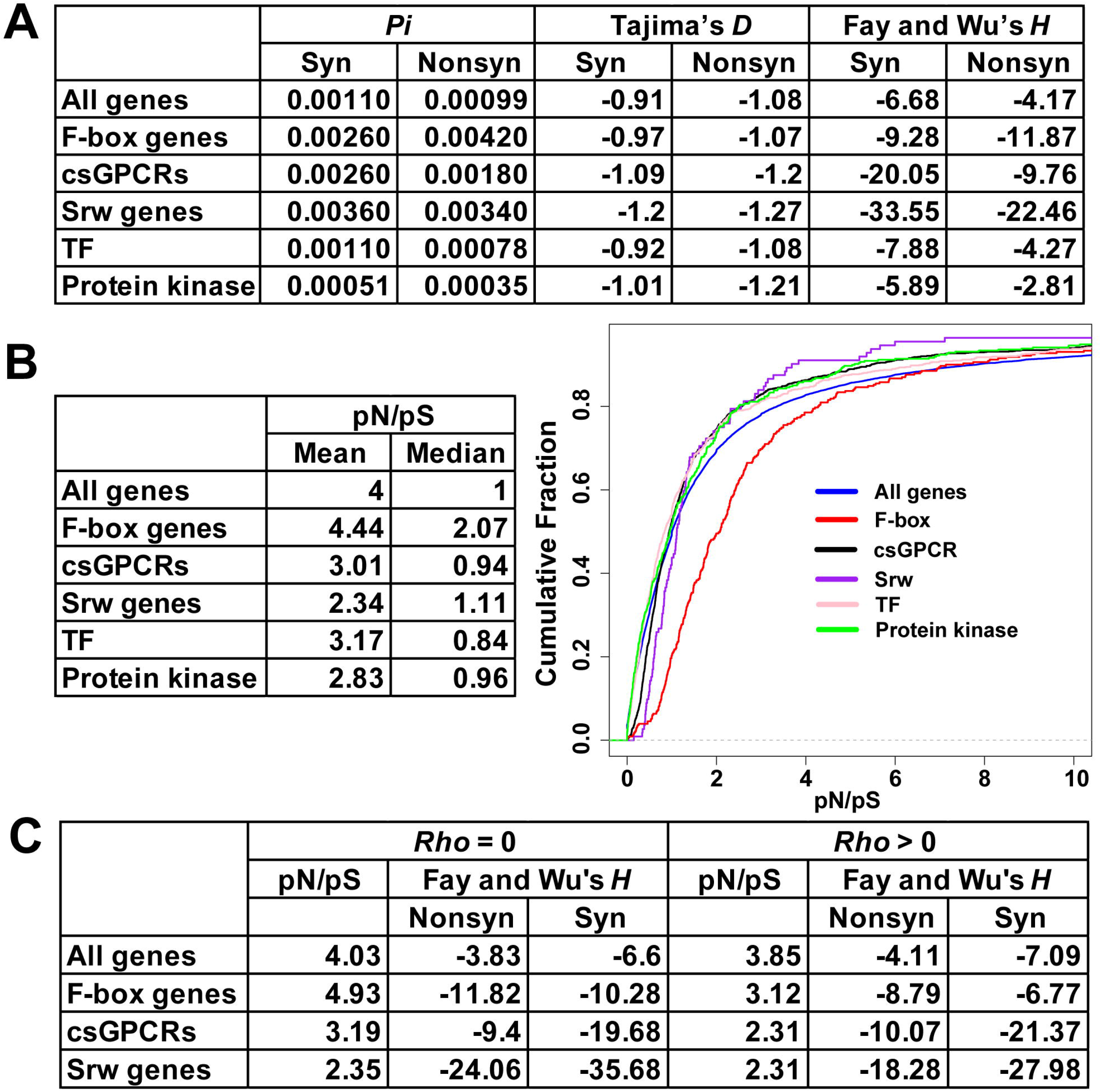
Selection on synonymous and nonsynonymous variants in F-box and csGPCR genes. (A) Mean values for *Pi*, Tajima’s *D*, and Fay and Wu’s *H* for different groups of genes calculated using synonymous or nonsynonymous SNVs. To compare the same set of genes for average *Pi*, we included the genes which has no synonymous or nonsynonymous SNVs (*Pi* = 0). So, the mean of *Pi* is slightly smaller than that in Figure 1C, which excluded the genes without nonsynonymous SNVs. (B) The mean and median of pN/pS ratios for different groups of genes and the cumulative distribution of the pN/pS ratios. (C) The average Fay and Wu’s *H* values for nonsynonymous and synonymous variants and the pN/pS ratios for different groups of genes in low and high recombination regions.

Interestingly, csGPCRs did not show higher than average pN/pS ratios and appeared to have a lot of synonymous SNVs, which have highly negative *H* values (fig. 6A). Some synonymous SNVs might be positively selected due to effects on codon usage and gene expression levels as previously seen in mammals (Resch, et al. 2007). Synonymous SNVs might also become high-frequency derived alleles through their linkage with positively selected nonsynonymous SNVs (Fay and Wu 2000). In fact, the *H* values for synonymous and nonsynonymous SNVs were highly positively correlated for F-box and csGPCR genes (supplementary fig. S8, Supplementary Material online).

F-box and csGPCR genes in low recombination regions appeared to be under stronger positive selection than genes in high recombination regions. We found that *H* values for both nonsynonymous and synonymous SNVs of F-box genes and *Srw* genes were more highly negative in low recombination regions than in high recombination regions (fig. 6C). In addition, pN/pS ratios were also higher in low recombination regions than in high recombination regions.

### Positive selection of F-box and csGPCR genes in non-Hawaiian population

The above analysis detected signs of strong positive selection in F-box and csGPCR genes among all wild strains. Among the populations, Fay and Wu’s *H* values were more highly negative in the non-Hawaiian strains than in the Hawaiian strains across all genes (fig. 5F). “Hawaii_3” group appeared to have lower *H* values than “Hawaii_1” and “Hawaii_2” groups probably due to the admixing with the non-Hawaiian strains. These observations are consistent with the selective sweep in non-Hawaiian populations. Genes in the F-box and csGPCR (especially *Srw*) genes showed much more highly negative *H* values than the genomic average not only in non-Hawaiian strains but also in Hawaiian strains, suggesting that they may also be under positive selection within the Hawaiian populations when considering XZ1516 as the most ancestral strain. Negative *H* values reflects the excess of high-frequency derived alleles. Being consistent with the above SNV analysis, the allele frequency of derived CNVs is much larger for F-box and csGPCR genes than the genomic average in both the entire population and the non-Hawaiian population of wild isolates, with XZ1516 as the outgroup (supplementary fig. S5B and C, Supplementary Material online).

Because XZ1516 is highly divergent, we also calculated *H* values for non-Hawaiian populations using a representative “Hawaii_1” (ECA396) or “Hawaii_2” (ECA742) strain as the outgroup. Similarly, F-box and csGPCR showed more highly negative *H* than the genomic average, indicating the accumulation of high-frequency derived alleles and possibly positive selection within the non-Hawaiian population, relative to the Hawaiian populations (fig. 5G).

### Different selection pressure on different domains of F-box and csGPCR proteins

F-box proteins all have two distinct funcitonal domains, a N-terminal F-box domain that mediates the assembly of SCF complex and a C-terminal substrate recognition domain that binds to the substrate proteins and targets them for ubiquitination. Using Pfam scan, we identified the F-box domain and putative substrate-binding domain (e.g. FTH, FBA, etc) in all F-box proteins and extracted the nonsynonymous SNVs mapped to these domains. Interestingly, *Pi* is much bigger and *H* much more negative for the SNVs mapped to the substrate-binding domain compared to those mapped to the F-box domain (fig. 7A). The enrichment of variants and stronger positive selection in the substrate-recognition domains supports the hypothesis that variations in the F-box genes may result in selective advantages by altering the ubiquitination and degradation of certain cellular proteins.

**Figure 7.**
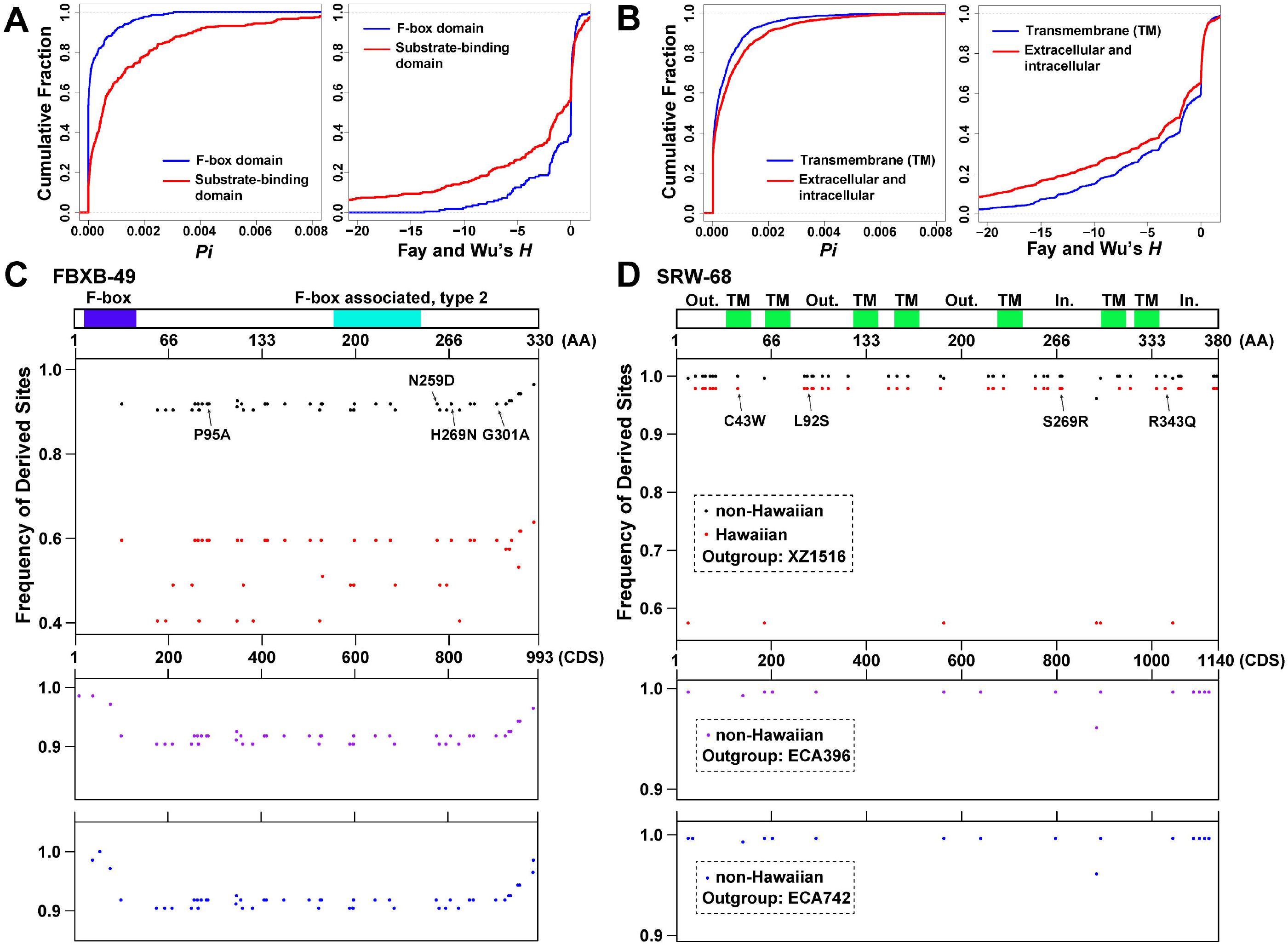
High-frequency derived sites were mapped to the substrate recognition domain of a reprehensive F-box protein and the extracellular loops of a representative csGPCR. (A) Cumulative distribution of *Pi* and Fay and Wu’s *H* for nonsynonymous SNVs in the F-box domain or putative substrate-binding domains of F-box proteins. (B) Distribution of *Pi* and *H* for SNVs in the transmembrane domain or extracellular or intracellular domains of csGPCRs. (C) The domain structure of a F-box protein encoded by *fbxb-49*. The F-box domain is in blue, and the type 2 F-box associated (FBA_2) domain, likely involved in binding substrate, is in cyan. (D) The domain structure of a csGPCR encoded by *srw-68*. The predicted transmembrane (TM) domain is in green. Extracellular loops (Out.) and intracellular (In.) tails are indicated. In both (A) and (B), the panel immediately below the domain structure indicate the position of high-frequency (>0.5) derived sites in non-Hawaiian populations using XZ1516 as the outgroup. Y-axis indicate the frequency of the derived alleles among the non-Hawaiian population (black dots) or the Hawaiian population (red dots). Each dot indicates a nonsynonymous SNVs. SNVs causing amino acid substitution with PROVEAN score below −2.5 were shown. The lower two panels showed the high-frequency derived sites in the non-Hawaiian population calculated using ECA396 (“Hawaii_1” strain; purple dots) or ECA742 (“Hawaii_2” strain; blue dots) as the outgroup.

As an example, F-box gene *fbxb-49* (*H* = −53.03) contains 48 high-frequency derived sites in the non-Hawaiian population (considering XZ1516 as the outgroup) and the frequency of those sites within the Hawaiian population are much lower (fig. 7C). Most of those sites are also high-frequency derived sites when using a “Hawaii_1” (ECA396) or “Hawaii_2” (ECA742) strain as the outgroup. Thus, selective sweep may have fixed those sites in the non-Hawaiian population. Very few variants are located in the region encoding F-box domain, while many more sites occurred in domains that are responsible for recognizing the substrate protein. Four of the sites have PROVEAN (Protein Variation Effect Analyzer) score (Choi and Chan 2015) below −2.5, suggesting potentially significant functional impacts.

Similarly, we mapped nonsynonymous SNVs onto the domain structure of csGPCRs and found that SNVs affecting the extracellular or intracellular regions of csGPCRs have larger *Pi* and more negative *H* than SNVs mapped to the transmembrane (TM) domains (fig. 7B). Conservation of amino acid sequences in the TM domain is expected, as the membrane protein topology may be maintained by purifying selection. Variations in the extracellular and intracellular regions could change the ability of the csGPCR to sense envrionmental cues and to transduce signals, respectively. So, these variants may be under positive selection and confer fitness advantages.

As an example, *srw-68* (*H* = −82.33) contains 44 high-frequency derived sites in non-Hawaiian population with XZ1516 as the outgroup. All of those sites are near fixation and mostly mapped to the extracellular or intracellular regions (fig. 7D). Four sites have PROVEAN scores below −2.5. Interestingly, many of these sites are also high-frequency derived sites in Hawaiian strains, suggesting that they are positively selected in the Hawaiian population as well. We also found 16 high-frequency derived sites in *srw-68* using the Hawaiian strains ECA396 and ECA742 as outgroup (fig. 7D), suggesting further divergence between the Hawaiian and non-Hawaiian alleles of this gene.

When comparing genes in low and high recombination regions, we found that the differences in the *Pi* and *H* values of nonsynonymous SNVs between domains of F-box proteins or csGPCRs persisted regardless of the recombination rate (supplementary fig. S9, Supplementary Material online).

### Extended Haplotype Homozygosity analysis identified selection footprints in F-box and csGPCR genes

In addition to the neutrality tests, we also applied the Extended Haplotype Homozygosity (EHH) method (Sabeti, et al. 2002) to detect the selection footprints among the nonsynonymous SNVs across the genome. EHH identifies long-range haplotypes and can discover genomic regions with strong selective sweep. First, we computed the integrated Haplotype Score (iHS) for both non-Hawaiian and Hawaiian strains (supplementary table S5, Supplementary Material online). Interestingly, the regions that showed extended haplotype homozygosity (high |iHS| scores) were in the left arms of chromosome II and III, where F-box genes are located, and the two arms of chromosome V, where most csGPCRs are located (fig. 8A and B). Indeed, among the 335 genes carrying at least one SNV with |iHS| > 2 in non-Hawaiian strains, csGPCR and F-box genes are enriched for 4.5 and 1.5 fold, respectively (fig. 8D), indicating that these genes may be under selective pressure among non-Hawaiian strains. Nevertheless, csGPCR and F-box genes may also be selected within the Hawaiian strains because of their enrichment in the regions with high |iHS| in the Hawaiian population (fig. 8D). This results is consistent with the highly negative Fay and Wu’s *H* for csGPCR and F-box genes in the Hawaiian population (fig. 5F).

**Figure 8.**
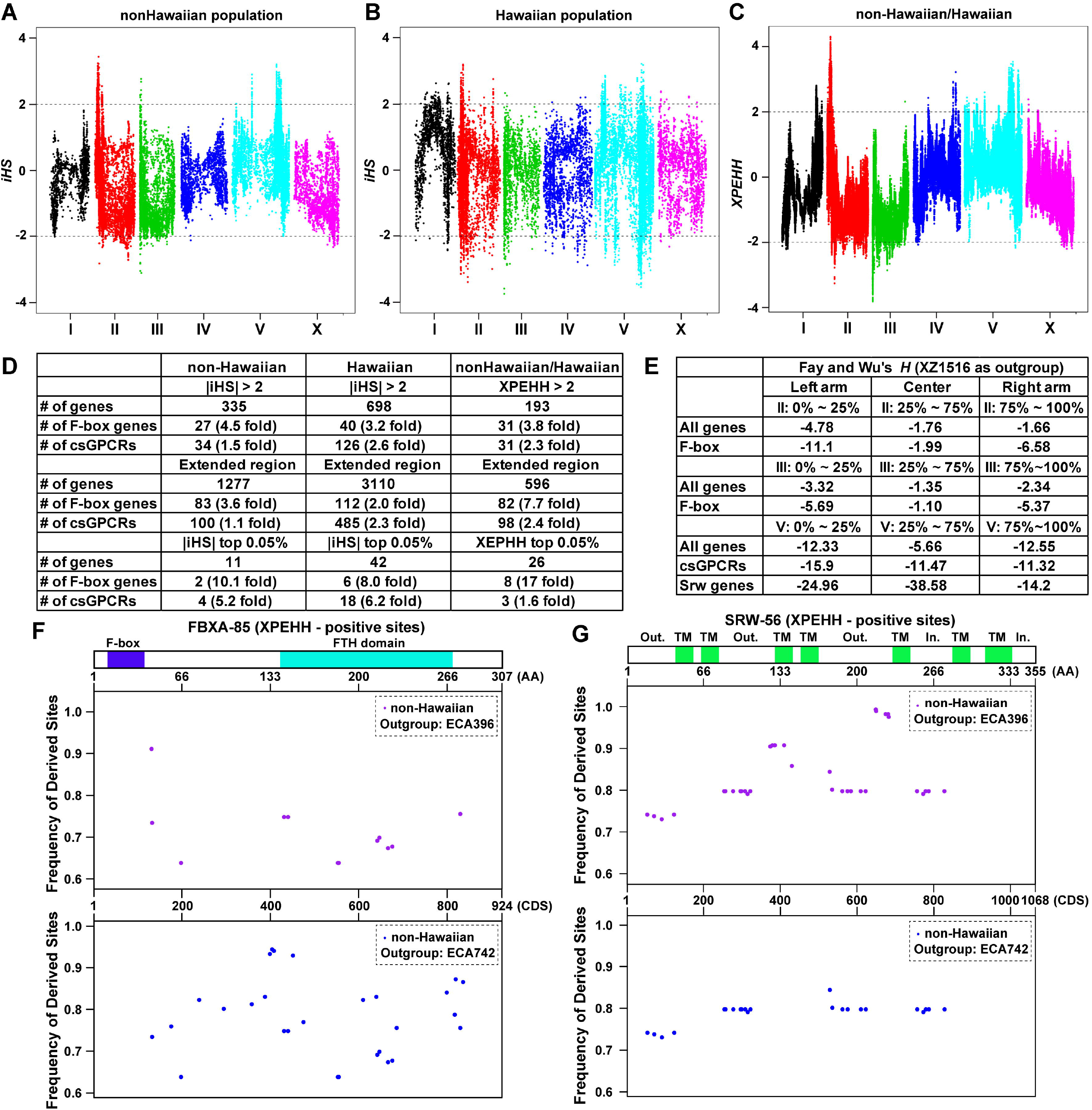
F-box and csGPCR genes are enriched in the genomic regions with selective footprint identified by extended haplotype homozygosity (EHH) analysis. (A-C) Manhattan plots of the extent of haplotype homozygosity measured by the integrated Haplotype Score (iHS) within the non-Hawaiian population (A) and Hawaiian population (B). (C) Regions of selection in non-Hawaiian population but not the Hawaiian population indicated by the Manhattan plots of cross-population EHH (XPEHH). (D) The number of F-box and csGPCR genes that contain SNVs with significant iHS or XPEHH and their folds of enrichment. For extended regions, significant SNVs that are less than 50-kb apart were connected to generate regions with selective footprints. (E) The mean Fay and Wu’s *H* values for all genes, F-box, and csGPCR genes in the arms and the center of chromosome (Chr) II, III, and V. (F) The domain structure of a representative F-box protein coded by *fbxa-85*; the F-box domain is in blue and the FTH domain in cyan. (G) The domain structure of a representative csGPCR coded by *srw-56*; the predicted transmembrane (TM) domain is in green, and extracellular loops (Out.) and intracellular (In.) tails are also indicated. Among the sites whose XPEHH > 2 in the two genes, the ones that are also high-frequency (> 0.5) derived sites with ECA396 (purple dots) and ECA742 (blue dots) as the outgroup are shown.

The genomic pattern of haplotype homozygosity is supported by the observations that *H* values of genes on the left arms of II and III are much more negative than the center and right arms and that *H* values of genes on both arms of V are smaller than the center of V (fig. 8E). F-box and csGPCR genes may be driving this pattern, because they tend to have even more negative *H* than average genes in the arms. In addition, Chromosome V generally had much more negative *H* than other chromosomes, suggesting signs of strong selective sweep, which is consistent with a previous observation of high haplotype homozygosity of V among non-Hawaiian strains (Andersen, et al. 2012). Selection of the over 1000 csGPCR genes on V may explain this chromosomal pattern.

Genes that are under selection in the non-Hawaiian population but not in the Hawaiian population may be associated with the adaptation of the non-Hawaiian strains. So, we conducted the XP-EHH (Cross-Population EHH) test to identify SNVs with such selection pattern and found the left arm of chromosome II and both arms of V contain regions with significantly positive XP-EHH values (fig. 8C). F-box and csGPCR genes are highly enriched in those regions. 18 out of the 41 genes carrying SNVs with XP-EHH > 2 on the left arm of II are F-box genes. The enrichment of F-box and csGPCR genes is even more obvious if we only consider the outlier SNVs (the top 0.05%) or count all genes in extended regions that connect significant SNVs within a 50-kb range (Mohd-Assaad, et al. 2018) (fig. 8D). In summary, both neutrality test and EHH analysis identified signs of strong positive selection on F-box and csGPCR genes in *C. elegans* wild isolates and especially in non-Hawaiian population.

As examples of highly selected genes, F-box gene *fbxa-85* carries 58 SNVs with significantly positive XP-EHH score (XP-EHH > 2; *p* < 0.05); 13 and 29 are high-frequency derived sites among non-Hawaiian strains using a “Hawaii_1” and “Hawaii_2” strain as the outgroup, respectively. Most of the sites occurred in the FTH domain involved in substrate binding and none in the F-box domain (fig. 8F). Similarly, csGPCR *srw-56* contains 67 SNVs with high XP-EHH; 36 and 24 of those SNVs are high-frequency derived sites in non-Hawaiian population with a “Hawaii_1” and “Hawaii_2” strain as the outgroup, respectively. Most of them occurred in the extracellular and intracellular domains of SRW-56 (fig. 8G).

### Selection patterns in F-box and csGPCR genes are not likely affected by varying population size and demographic history

Varying population size and demographic history are known confounding factors for predicting selective sweep (Wakeley and Aliacar 2001; Przeworski 2002; Nielsen, et al. 2005). We next addressed whether these two factors confounded our neutrality test results. To assess whether the varying number of strains in the 11 subpopulation among the wild isolates had effects on the neutrality test statistics, we selected strains and SNVs using two different sampling schemes (scattering and pooling schemes) according to previous studies (Stadler, et al. 2009; Li, et al. 2010) (see Materials and Methods). Polymorphism, Tajima’s *D*, and Fay and Wu’s *H* calculated using the samples obtained with the two sampling methods are very similar (supplementary fig. S10, Supplementary Material online), indicating that the varying population sizes among the subpopulation do not significantly confound our results.

Previous population history analysis of *C. elegans* found that wild isolates in non-Hawaiian population may have suffered a strong decline in population size about 10,000 generations ago (Thomas, et al. 2015). To assess the confounding effect of the potential bottleneck on selection detection, we simulated SNV data under neutrally constant population size model or bottleneck model and plotted site frequency spectra (SFS). Bottleneck leads to the enrichment of low and high-frequency alleles in simulated data as expected (supplementary fig. S11, Supplementary Material online). However, SFS pattern of the empirical SNV data of non-Hawaiian populations are more similar to the constant population size model, suggesting that the potential bottleneck effect may not significantly change the site frequency in the *C. elegans* wild isolates we analyzed.

We next predicted selective sweep sites based on the site frequency spectrum using the software SweeD, which analyzes composite likelihood and is robust against recombination and demographic assumption (Nielsen, et al. 2005). We found that selected sites at the significance threshold of 1% are mostly located in the arms of chromosomes (supplementary fig. S12, Supplementary Material online), where F-box and csGPCRs are enriched, which is consistent with the results of neutrality tests and EHH analysis. Among the 564 significant sites located in 233 genes (mean α score, an indicator of selection coefficient, is 31), 31 sites are mapped to 10 F-box genes (2.5 fold enrichment) with an average α score at 54. csGPCR genes carry 28 significant sites with an average α score at 63. Thus, even considering demographic history, F-box and csGPCR genes still show strong positive selection.

## Discussion

The nematode *C. elegans*, which is traditionally used as a model organism for molecular biology, has emerged as an important organism in studying the genetic mechanisms of evolution. The genomic sequences of over 50 species in the *Caenorhabditis* genus and 330 wild *C. elegans* isotypes provided an important resource for understanding the evolutionary history of *C. elegans* and nematodes in general, e.g. the rise of self-fertile hermaphroditism through convergent evolution in *C. elegans* and *C. briggsae* (Nayak, et al. 2005) and the balancing selection maintaining genetic incompatibilities among *C. elegans* wild isolates (Seidel, et al. 2008). In this study, we aimed to identify genes or gene families that have large diversity among the *C. elegans* wild isolates and show signs of positive selection. The F-box gene family and chemosensory GPCR genes emerged from our analysis, suggesting that they may contribute to the adaptation of wild *C. elegans*.

### Intraspecific positive selection of F-box genes

Compared to insects and vertebrates, *C. elegans* genome contains a large number of F-box genes. This increased number of F-box genes might have allowed selective recognition of target proteins for degradation in a precisely controlled manner and the increased precision in the regulation of protein turnover might have contributed to nematode evolution. In fact, an earlier study calculated the nonsynonymous (*dN*)/synonymous (*dS*) ratio among paralogous F-box genes in *C. elegans* reference genome (the N2 strain) and found evidence of purifying selection in the sequence encoding the F-box domain and positive selection in the substrate recognition domain (Thomas 2006). Our studies using the genomic sequences of 330 *C. elegans* wild isolates found large intraspecific variations in the F-box genes and signs of strong positive selection, which may imply their roles in adaptation. Interestingly, variants in the substrate-binding domain showed larger polymorphism and stronger selection than the variants in the F-box domain, supporting that the function of substrate recognition but not Skp1 binding is the target of positive selection.

What kind of selective advantages can variants in F-box genes confer? Recent studies suggested a link between the SCF complex and antimicrobial immunity in *C. elegans*, because the transcription of many components of the SCF complex were upregulated upon Orsay virus and *Nematocida parisii* (a microsporidia fungi) infections (Chen, et al. 2017) and RNAi knockdown of the core SCF components promoted the infection (Bakowski, et al. 2014). Among the upregulated genes are F-box genes that show strong signs of positive selection in our studies, e.g. *fbxc-19, fbxa-75, fbxa-135, fbxa-158, fbxa-165*, and *fbxa-182*, whose Fay and Wu’s *H* are all below −20. Thus, an attractive hypothesis is that variations in F-box proteins allow or enhance the ability of SCF complex to ubiquitinate microbial and/or host proteins required for the replication of the pathogen, thus contributing to stronger immune defence. In addition to antiviral immunity, we also expect certain alleles of F-box genes to confer other fitness advantages, given the importance of ubiquitination-proteasome system in many biological processes.

### Intraspecific adaptive evolution of csGPCRs

The csGPCR family is the largest gene family in *C. elegans* and contains over 1,300 genes. Through the studies of specific phenotypes, a few csGPCRs were previously connected to adaptation (Dennis, et al. 2018; Lee, et al. 2019). For example, the deletion of two csGPCR genes, *srg-36* and *srg-37*, which resulted in insensitivity to the dauer pheromone ascaroside and defects in entering dauer diapause, were acquired independently by two domesticated *C. elegans* strains grown in high density (McGrath, et al. 2011). Similar loss-of-function deletions in *srg-36* and *srg-37* were also found in natural isolates across the globe, suggesting that niche-associated variation in pheromone receptors may contribute to the boom-and-bust population dynamics (Lee, et al. 2019). In addition, a frameshift-causing deletion in another csGPCR, *str-217*, in Hawaiian strain CB4856, led to resistance to the insect repellents N,N-Diethyl-meta-toluamide (DEET) (Dennis, et al. 2018) and similar deletions were found in nine other wild isotypes (our unpublished results), suggesting that *C. elegans* may have evolved to acquire resistance to harmful environmental chemicals by inactivating csGPCRs. The above examples showcased how the intraspecific evolution of individual csGPCR genes can have significant functional consequences and contribute to adaptation. Our study, in a more systematic way, indicates that csGPCRs are highly diverse and are under strong positive selection in the *C. elegans* wild population.

Among the four csGPCR superfamilies (*Str, Sra, Srg*, and *Solo*), our analysis using nonsynonymous SNVs found that *Str* genes had larger polymorphism and stronger positive selection than *Sra* and *Srg* genes (fig. 1C and 5D), which is consistent with previous observation on the intraspecific variations of *Str* genes (Stewart, et al. 2005). In fact, these variations created ∼200 pseudogenes *Str* genes in *C. elegans* reference genome (the N2 strain) through often times only one apparent defect. Compared to *Str* genes, we found that *Solo* superfamily csGPCRs, especially *Srw* genes have even larger diversity and stronger positive selection. Interestingly, *Srw* genes appeared to be more ancestral than the other csGPCR families and likely originated from the large Rhodopsin GPCR family before the split of the nematode lineage (Krishnan, et al. 2014). *Srw* genes are the only csGPCRs that have clear homology with insect and vertebrate GPCRs and likely code for FMRFamide/peptide receptors (Robertson and Thomas 2006), so variations in these genes might lead to selective advantages in peptide sensing. Moreover, most of the high-frequency derived sites of *Srw* genes are mapped to the regions that code for the extracellular and intracellular domains, suggesting that altered ligand recognition and/or signal transduction might be positively selected. A similar observation was made for *Srz* genes in the *Solo* superfamily based on that *dN/dS* ratios among paralogous groups of *Srz* genes in *C. elegans* and *C. briggsae* peak in the extracellular loops (Thomas, et al. 2005). Thus, the large gene pool of csGPCRs may facilitate the adaptation to a changing environment by supplying alleles with specific ligand-binding or signalling properties for positive selection.

### The correlation between large diversity and strong positive selection

Compared to other gene families, F-box and csGPCR genes not only have large genetic diversity but also show strong signs of positive selection. We reason that gene families such as the TFs and protein kinases have low polymorphism because they play critical roles in the development of *C. elegans* and thus may be under purifying selection. In comparison, F-box and csGPCR genes maintain large polymorphism likely due to the lack of strong purifying selection, as well as high recombination rate and frequent gene flow. High recombination rate results from their clustering in the chromosomal arms, and gene flow between genetically divergent subpopulations helps maintain genetic diversity.

Rapid expansion of the F-box and csGPCR gene families and high rate of gene gain and loss also contributed to their large diversity and facilitated positive selection and adaptation. Indeed, our analysis of copy number variants found more frequent gene duplication and deletion in F-box and csGPCR genes than genomic average, supporting fast intraspecific evolution of these genes. Previous studies found that the *C. elegans* genome shows a higher duplication rate than *Drosophila* and *yeast* genomes (Lipinski, et al. 2011). This pattern is likely driven by the duplication of F-box and csGPCR genes. Functional diversification of the duplicated genes could lead to novel functional characteristics. Although the function of most F-box and csGPCR genes in *C. elegans* are unknown, their expression pattern, to certain extent, reflects their potential functions. For example, among the 39 positively selected (H < −20) csGPCRs whose expression were studied before (Vidal, et al. 2018), we found that these csGPCRs show distinct expression patterns in a diverse range of tissues (supplementary fig. S13, Supplementary Material online). Although expression is heavily enriched in sensory neurons, most csGPCRs are expressed in unique sets of cells and identical expression patterns for two csGPCRs are rare. We suspect the diversification in expression regulation is correlated with diversification in functions. Thus, our data supports a model that duplications of F-box and csGPCR genes and accumulation of nonsynonymous SNVs lead to functional diversities in protein degradation and chemosensation pathways, which allowed positive selection to act upon.

## Materials and Methods

### Population genetic statistics

To obtain the genomic data of *C. elegans* wild isolates, We used the hard-filtered VCF (Variant Call Format) file (20180527 release) provided by the *C. elegans* natural diversity resource (CeNDR; https://www.elegansvariation.org/) (Cook, et al. 2017). We chose the hard-filtered VCF over the soft-filtered VCF to avoid including low-quality reads and variants with low coverage depth in our analysis. The hard-filtered VCF file contained in total 2,906,135 high-quality variants, including 2,493,687 SNVs and 412,448 small indels, which were annotated by SnpEff (v4.3t) using the Ensemble WBcel235.94 genome assembly. About half (1,124,958) of the SNVs were found in only one of the 330 isotypes, and they all occurred as homozygotes likely due to the hermaphroditism-driven homozygosity in *C. elegans*; we consider those SNVs as singleton (or private doubleton) and included them in most of our analysis. Among the 2,906,135 variants, 594,265 occurred in the protein-coding region (CDS) and 2,311,870 occurred in non-coding regions. 266,004 SNVs caused nonsynonymous mutations, 271,718 SNVs caused synonymous mutations, and 51,701 SNVs may affect mRNA splicing.

Among all SNV sites, we found that 665,368 SNVs in 11,199 genes had complete sequencing data in all 330 wild strains (660 alleles) using VCFtools (v0.1.13) (Danecek, et al. 2011). This dataset is referred to as “the complete-case dataset”. SNVs in the complete-case dataset were then subjected to calculation using DnaSP (Rozas, et al. 2017) and PopGenome (Pfeifer, et al. 2014). Both software produced similar results for nucleotide diversity (*Pi*) and neutrality test statistic Tajima’s *D* for the synonymous, nonsynonymous, intron and UTR sites (supplementary fig. S1, Supplementary Material online). Correlation analysis was done in R (v3.6.1) using Pearson correlation test (R function cor.test).

Because the analysis of the complete-case dataset removed 73% of the variant sites, we tested whether similar results can be obtained if we include variant sites with incomplete data. For an average strain, 76,872 (2.6%) out of the total 2,906,135 variant sites did not have high-quality sequencing data. For a variant site, 17.5 (2.6%) on average (median value is 3.9 (0.6%)) out of the 660 alleles (330 strains) did not have valid genotypes. Stringent analyses using only complete datasets discarded almost three quarters of the SNVs and possibly lost valuable information. To deal with this problem, we used the software VariScan (Hutter, et al. 2006) to set a threshold for the number of alleles containing valid data for a given site. We first annotated the VCF to extract nonsynonymous SNVs and converted the VCF formatted file to Hapmap style using Tassel (v5.0) (Bradbury, et al. 2007) to facilitate the calculation of *Pi* (Nei 1987), Tajima’s *D* (Tajima 1989), and Fay and Wu’s *H* (Fay and Wu 2000) by VariScan. We then tested the threshold (NumNuc) at 200 and 450, where the sites with more than 200 and 450 alleles were used for the analyses, respectively. These two conditions included 253,600 and 235,283 nonsynonymous variants covering 18,797 and 18,643 genes, respectively, as compared to the complete-case dataset that contained only 85,260 sites covering only 9,948 genes. Preserving more SNVs lead to larger *Pi*, while Tajima’s *D* and Fay and Wu’s *H* did not change (supplementary fig. S1, Supplementary Material online). Thus, the inclusion of variant sites with a few missing data points did not affect the results of neutrality test, but a significant amount of genetic diversity data were kept. For most analyses, we opted to use the dataset that included all sites with >200 valid alleles. This dataset is referred to as “the full dataset”.

To assess the significance of the *D* and *H* values, we performed coalescent simulations (Hudson 1990; Librado and Rozas 2009) for each gene based on the number of segregating sites using DnaSP v5. The confidence interval was set as 95% and the number of replicates was 1000. We found that vast majority (>95%) of the *D* value smaller than −2 and *H* value smaller than −20 have *p* values lower than 0.05.

### Population structure analysis

We first used PLINK (v1.9) (Purcell, et al. 2007) to convert the VCF file containing 2,493,687 SNVs to a PED formatted file, which was then subjected to the analysis using ADMIXTURE with the number of subpopulation (K value) ranging from 2 to 15. The cross-validation (CV) error for K=11 was the smallest. The population structure was visualized using the pophelper web apps (v1.0.10) (Francis 2017). The 11 ancestral groups are: Europe_1, Europe_2, Europe_3, Europe_4, Europe_5, Europe_6, Hawaii_1, Hawaii_2, Hawaii_3, North America and Australasia, which were named based on the geographic locations of most strains that carry the ancestral lineage (supplementary fig. S2 and table S1, Supplementary Material online). Some correlation between geographical separation and genetic divergence were seen, e.g. “Europe_5” and “Europe_6” strains were mostly found on Iberian Peninsula and Portuguese islands. Out of the 330 strains, 266 have one dominant ancestral lineage (one ancestral proportion > 0.5); the other 64 strains showed considerable mixing between at least three ancestral populations. “Hawaii_1” and “Hawaii_2” are the same as the previous “Hawaiian Volcano” and “Hawaiian Divergent” subpopulations, and “Hawaii_3” is a combination of the “Hawaiian Low” and “Hawaiian Invaded” subpopulations defined by Crombie, et al. (2019).

We then grouped the 330 wild isolates into Hawaiian and non-Hawaiian populations based on genetic difference instead of geographic locations (supplementary table S2, Supplementary Material online). The Hawaiian population contains 45 strains carrying a dominant lineage (admixing proportion > 0.5) from “Hawaii_1” (10 strains), “Hawaii_2” (10 strains), and “Hawaii_3” (25 strains). The remaining 285 strains were grouped as the non-Hawaiian population, among which sixty-four strains did not have a dominant ancestral lineage and contained extensive admixing among mostly the eight non-Hawaiian ancestral subpopulations. They were, thus, included in the non-Hawaiian population. The 45 strains in the Hawaiian population were all extracted from Hawaiian Islands except five strains (ECA36, JU3226, QX1211, ECA593, and XZ2019), and five strains that were extracted from Hawaiian Islands were included in the non-Hawaiian population (ECA928, ECA923, ECA369, QX1791, and XZ1515) because they were genetically very different from Hawaiian strains.

This grouping of Hawaiian and non-Hawaiian populations was used for the computation of polymorphism (*Pi*), Tajima’s *D*, and Fay and Wu’s *H* within each population and were used for extended haplotype homozygosity (EHH) analysis (described later). For the calculation of F_ST_ and the gene flow and migration analysis among the 11 subgroups, we removed the strains without any ancestral proportion over 0.5 and kept 221 strains for the eight non-Hawaiian subpopulations and 45 strains for the three Hawaiian subpopulations.

### Phylogenetic analysis

To visualize the phylogenetic relationship of the Hawaiian and non-Hawaiian populations, we used nonsynonymous SNVs from all 45 Hawaiian strains and 24 non-Hawaiian strains (3 strains with the biggest ancestral proportion from each subgroup). These 24 strains represented the genetic diversity of the non-Hawaiian population, allowing easy visualization without making the tree too crowded. We used Tassel to convert VCF file to Phylip interleaved format and constructed the neighbour-joining net with SplitsTree4 (v4.15.1) (Huson and Bryant 2006). For the trees with just csGPCR and F-box genes, we used VCFtools to extract nonsynonymous SNVs of these genes according to their genomic location. When making the tree by SplitsTree, one thousand bootstrap replicates were performed. Edges with 100% bootstrap support are labelled with “100”.

*Caenorhabditis briggsae, Caenorhabditis remanei*, and *Caenorhabditis brenneri* were chosen as the outgroups. The coding sequences of *C. elegans* genes and their orthologs in *C. briggsae, C. remanei*, and *C. brenneri* were downloaded from WormBase (WS275) and then aligned using MegaX (Kumar, et al. 2018) to identify variants. We used a set of algorithms, including OrthoMCL, OMA, TreeFam, ParaSite-Compara, Inparanoid_8, WormBase-Compara, and Hillier-set, and merged the results to identify the orthologs of *C. elegans* genes in the other three species. We then checked each nonsynonymous SNV that existed in the *C. elegans* wild isolates (VCF file from CeNDR) for their presence in *C. briggsae, C. remanei*, and *C. brenneri* genomes. If the allele in the three species matched the *C. elegans* reference (N2) sequence, it was considered as a wild-type; if the allele matches the alternative sequence, the species carried that variant. If neither, we considered it missing for that SNV. In the case of one species having multiple orthologs of the *C. elegans* genes, we checked the SNV against all orthologs and if any of them had the alternative sequence, we considered the species to carry the variant. In total, we found 78,833, 74,274, and 55,234 *C. elegans* SNVs in *C. briggsae, C. remanei*, and *C. brenneri*, respectively and included these data in the tree reconstruction.

### Gene family analysis and gene enrichment analysis

Based on previous publications, we compiled a list of genes in the csGPCR gene family (Vidal, et al. 2018), F-box gene family (Kipreos and Pagano 2000; Thomas 2006), transcription factor family (Reece-Hoyes, et al. 2005), and protein kinase family (Manning 2005). For tissue-specific genes, we collected genes whose expression are enriched in muscle, intestine, germline (Pauli, et al. 2006), and neurons (Von Stetina, et al. 2007). To compare *Pi*, Tajima’s *D*, and Fay and Wu’s *H* values between different groups of genes, we performed non-parametric Wilcoxon’s test to evaluate the statistical significance of the difference between groups.

For gene enrichment analysis, simple enrichment fold of csGPCR and F-box genes are calculated as observed gene frequency divided by expected gene frequency. We also subjected a list of specific genes to Gene Set Enrichment Analysis at wormbase.org (Angeles-Albores, et al. 2018). Q value threshold cutoff was set at 0.1 to generate results of Tissue Enrichment Analysis (TEA), Phenotype Enrichment Analysis (PEA), and Gene Enrichment Analysis (GEA).

### Fixation index (F_ST_) calculation

Hudson’s F_ST_ (Hudson, et al. 1992) were estimated using PopGenome. SNVs from the 266 strains that have an ancestral proportion bigger than 0.5 (221 non-Hawaiian and 45 Hawaiian strains) were subjected to the calculation. Prior to computation, we removed SNVs with valid genotype data in less than 100 strains to be consistent with VariScan analysis (NumNuc = 200).

### Gene flow analysis

The migration events among subpopulations were analyzed by TreeMix (Pickrell and Pritchard 2012). We first used Stacks (Catchen, et al. 2013) to convert VCF file into the input format required by treemix. In each run, 1000 SNP blocks were set for all genes, and 100 SNP blocks were set for the analysis of csGPCR or F-box genes. Hawaii_1 was used as the outgroup and three migration events were allowed. On GNU parallel, one thousand bootstrap replicates were performed for all five analyses. From the bootstrap results, we extracted the common migration events and calculated the probability of occurrence for each migration events among the 1000 replicates. The top three events were presented. We also calculated the average migration weight for each of the three events among the 1000 bootstrap replicates and the average weight were color-coded. To avoid possible interference by singletons and linkage disequilibrium, we repeated the analysis after removing the singletons and highly linked SNPs (using plink --indep-pairwise 50 10 0.8) and obtained very similar results.

### Estimation of recombination rate

Recombination rates were estimated using all 2,493,687 SNVs and the R package, FastEPRR (Gao, et al. 2016). We set the sliding window size to 50,000 bp and the sliding step length to 25,000 bp. After obtaining the estimated recombination rate for each genomic window, we assigned that recombination rate (*Rho* value) to genes, whose CDS range overlaps with the genomic window.

### Extended haplotype homozygosity (EHH) analysis

We used EHH analysis to identify regions with selection footprints (Sabeti, et al. 2002). VCF file was first phased by beagle (v5.1) (Browning and Browning 2007) and then subjected to haplotype analysis using the rehh (v3.0) R package (Gautier and Vitalis 2012) to calculate the Integrated Haplotype Score (iHS) and the Cross-population Extended Haplotype Homozygosity (XPEHH) value. Strains were grouped as non-Hawaiian and Hawaiian as described above when computing iHS and XPEHH. Unpolarized data were used to avoid making assumption of ancestry.

### Assessing the influence of varying population size and bottleneck effects

Because different subpopulations have different numbers of strains, the varying population size may create bias when calculating neutrality statistics. We assessed this potential bias by comparing the SNV data extracted through different sampling schemes (scattering and pooling schemes) as previously established (Stadler, et al. 2009; Li, et al. 2010). For scattered sampling, we randomly selected 5 strains from each of the 11 subpopulations based on population structure; for pooled sampling, we randomly selected 15 Hawaiian strains (3 subpopulations) and 40 non-Hawaiian strains (8 subpopulations). We repeated the sampling 100 times and then calculated the average *Pi*, Tajima’s *D*, and Fay and Wu’s *H*. The small differences in their values between the scattered and pooled sampling schemes suggest that the bias introduced by varying population size is not significant.

To assess the influence of demographic history and bottleneck effects on the neutrality tests, we simulated SNV data using the software MSMS (Ewing and Hermisson 2010) under constant population size model and bottleneck model. Parameters for the simulation were set according to previous studies (Andersen, et al. 2012). The command for simulating the two models are: msms -N 20000 -ms 440 1000 -t 100 -r 150 -SAA 500 -Sp 0.5 -SAa 200 (constant) and msms -N 20000 -ms 440 1000 -t 100 -r 150 -SAA 500 -Sp 0.5 -SAa 200 -eN 0.015 0.01 -eN 0.020 1.0 (bottleneck). The simulated data were then plotted as site frequency spectra (SFS), which were compared to the empirical site frequency spectrum data for nonsynonymous SNVs in *C*.*elegans*.

We also used the software SweeD to estimate the selective sweep position. SweeD appeared to be robust against the confounding effect of bottleneck on selective sweep prediction (Nielsen, et al. 2005; Pavlidis, et al. 2013). We identified the selected sites with significant score using the likelihood threshold of 0.01. Genes that harbour these selected sites were then identified.

### Copy number variation (CNV) analysis

The raw sequencing data of the 330 wild isolates were downloaded from NCBI (PRJNA549503). Sequencing reads were aligned to the reference genome of *C. elegans* using BWA-mem (v0.7.17). Structural variants were called using Manta (Chen, et al. 2016). The output VCF was merged by bcftools (v1.9). Structural variants with <= 5bp position difference and <=20% size difference were merged together. Deletions and duplications were considered as copy number variation. Large deletions or duplications with more than 1 Mbp and chromosome-level variation were discarded. Derived CNV allele frequency were calculated using XZ1516 as the outgroup.

### Protein domain structure and PROVEAN score

We used PfamScan tools (ftp://ftp.ebi.ac.uk/pub/databases/Pfam/Tools/) to identify the F-box domain and the potential substrate-recognition domains (e.g. FTH, FBA, HTH-48, WD40 repeats, LRR, etc) of F-box proteins. We used the TMHMM server (http://www.cbs.dtu.dk/services/TMHMM/) to predict the transmembrane domain (TM) and the intracellular and extracellular regions of csGPCR proteins. SNVs falling into these different domains were then filtered accordingly. The potential functional effect of the nonsynonymous mutations is predicted by the PROVEAN (Choi and Chan 2015) web server (PROVEAN v1.1.3) at http://provean.jcvi.org/index.php; a score lower than −2.5 is deemed as having significant effects on the function.

## Acknowledgments

We thank the *Caenorhabditis elegans* Natural Diversity Resource (CeNDR) at Northwestern University for sharing genomic data of the *C. elegans* wild isolates through their website. This study is supported by funds from the Research Grant Council of Hong Kong [ECS 27104219], the Food and Health Bureau of Hong Kong [HMRF 07183186], and seed funds from the University of Hong Kong [201812159005 and 201910159087] to C.Z. Computational work were performed using research computing facilities offered by Information Technology Services at the University of Hong Kong.

## Data availability

VCF files and raw sequencing data used in this study can be downloaded from the CeNDR website (https://www.elegansvariation.org/). Most of our computational results can be found in supplemental tables. Other data, e.g. the simulated SNV data and the VCF file for CNVs, are available from the authors upon reasonable request.

