## Supplemental Figures for "Large genetic diversity and strong positive selection in F-box and GPCR genes among the wild isolates of *Caenorhabditis elegans*"

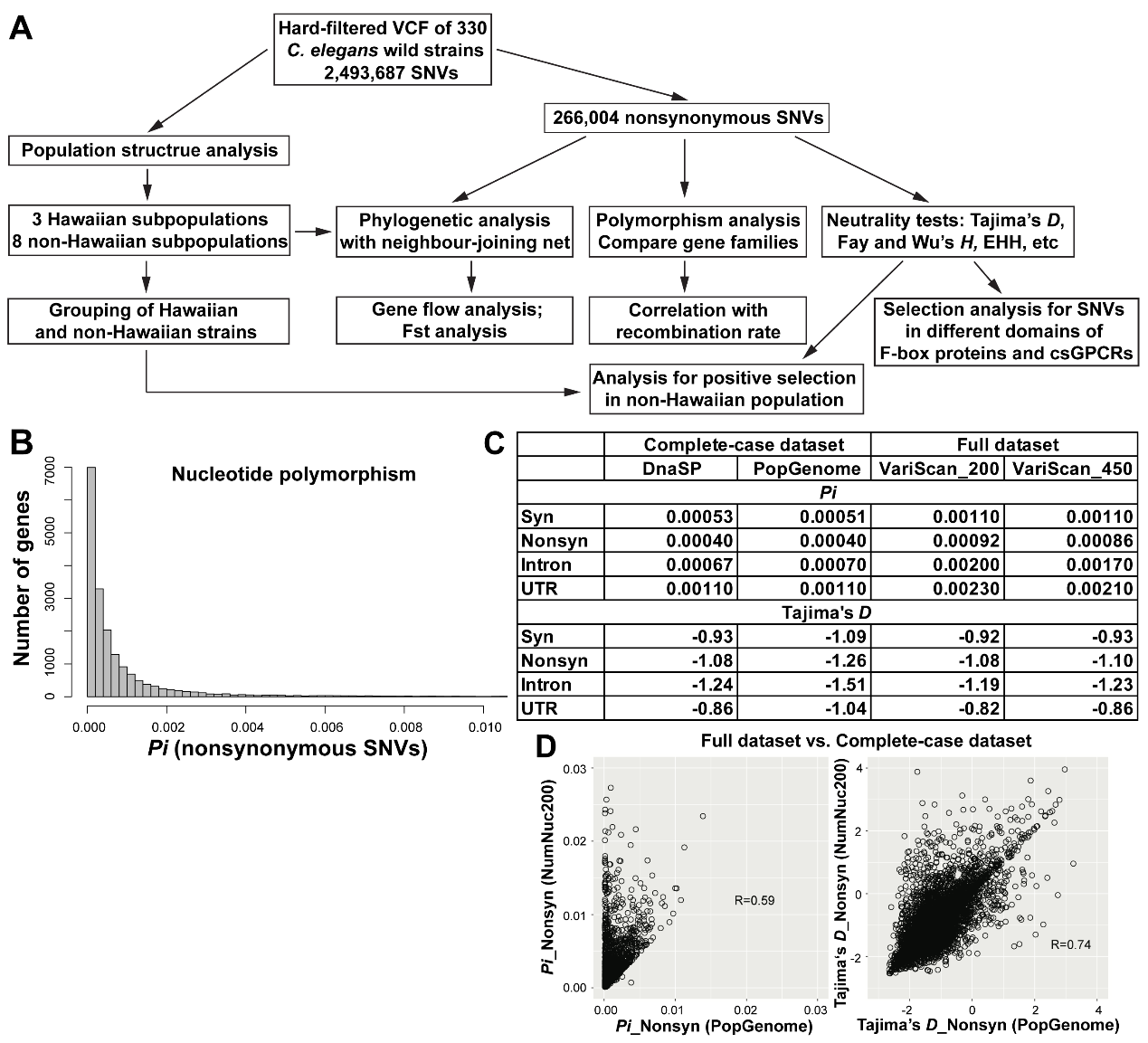

**Supplementary Figure S1. Flowchart of the analysis for the polymorphism and neutrality test across all genes.** (A) Flowchart of the analysis of SNVs among the wild isolates of *C. elegans*. (B) A histogram for the distribution of *Pi* across all genes. *Pi* values of individual genes can be found in Table S2. (C) The mean of *Pi* and Tajima’s *D* calculated for synonymous, nonsynonymous, intron and UTR SNVs in all genes using different software. (D) The correlation of *Pi* and Tajima’s *D* computed using the full dataset (VariScan; NumNuc = 200) and the complete-case dataset (PopGenome; excluding sites with any missing genotype data).

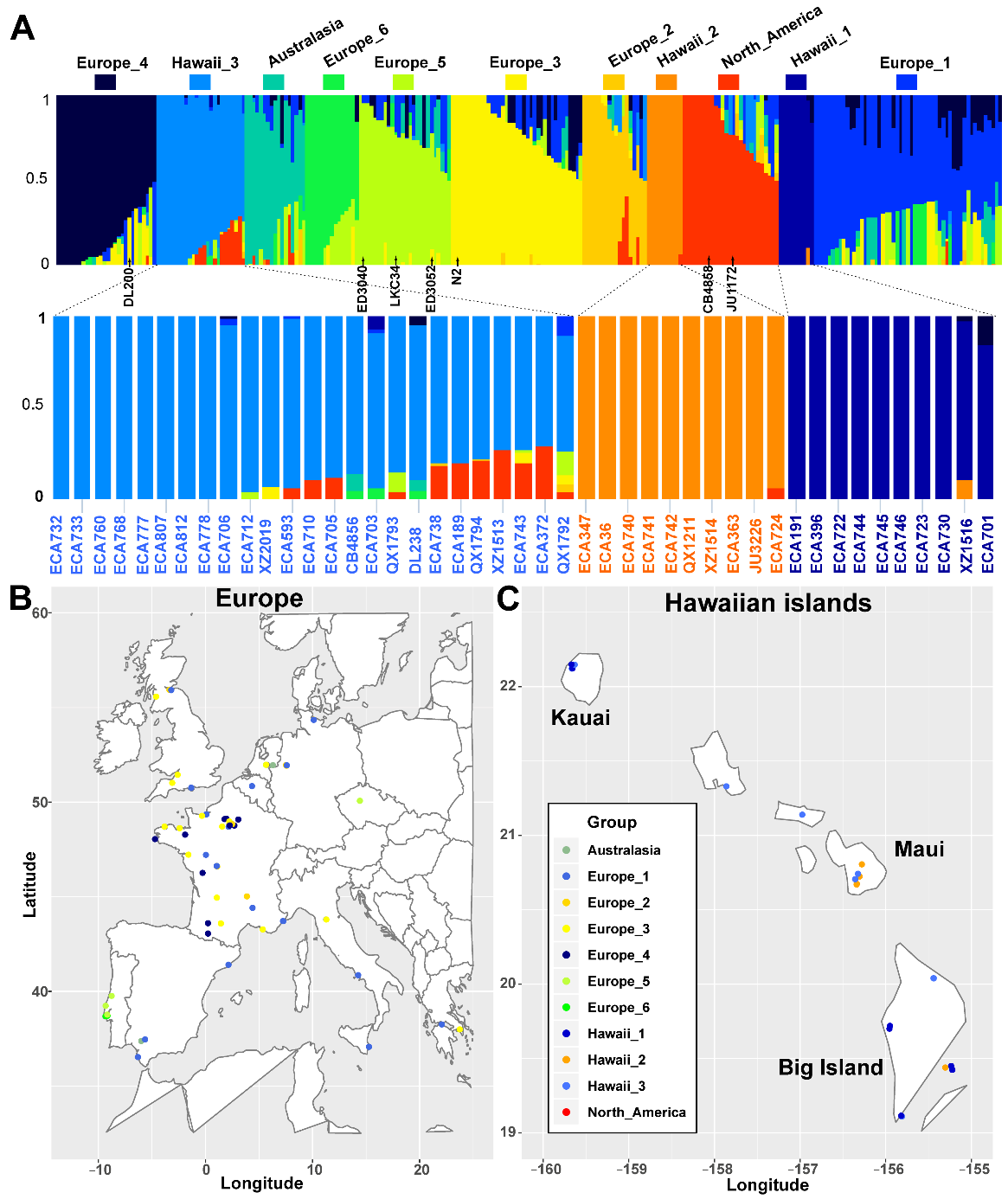

**Supplementary Figure S2. Population structure and geographical distribution of *C. elegans* wild isolates.** (A) The ancestral population proportions of wild isolates (color-coded) inferred by Admixture, when the number of population (K value) was set at 11. The bar indicates the ancestral fraction for each strain. Only strains with one ancestral proportion larger than 0.5 were shown. Each group (or subpopulation) was named after the geographical location of most strains in that group. The names of the strains from the three Hawaiian subpopulations were shown in the enlarged panels. (B-C) Geographical locations of strains isolated in Europe (B) and Hawaii (C). Each dot represents a strain, but some dots are entirely overlapping given the scale of the map. The latitude and longitude data for each strain was obtained from CeNDR. Color code is consistent with (A).

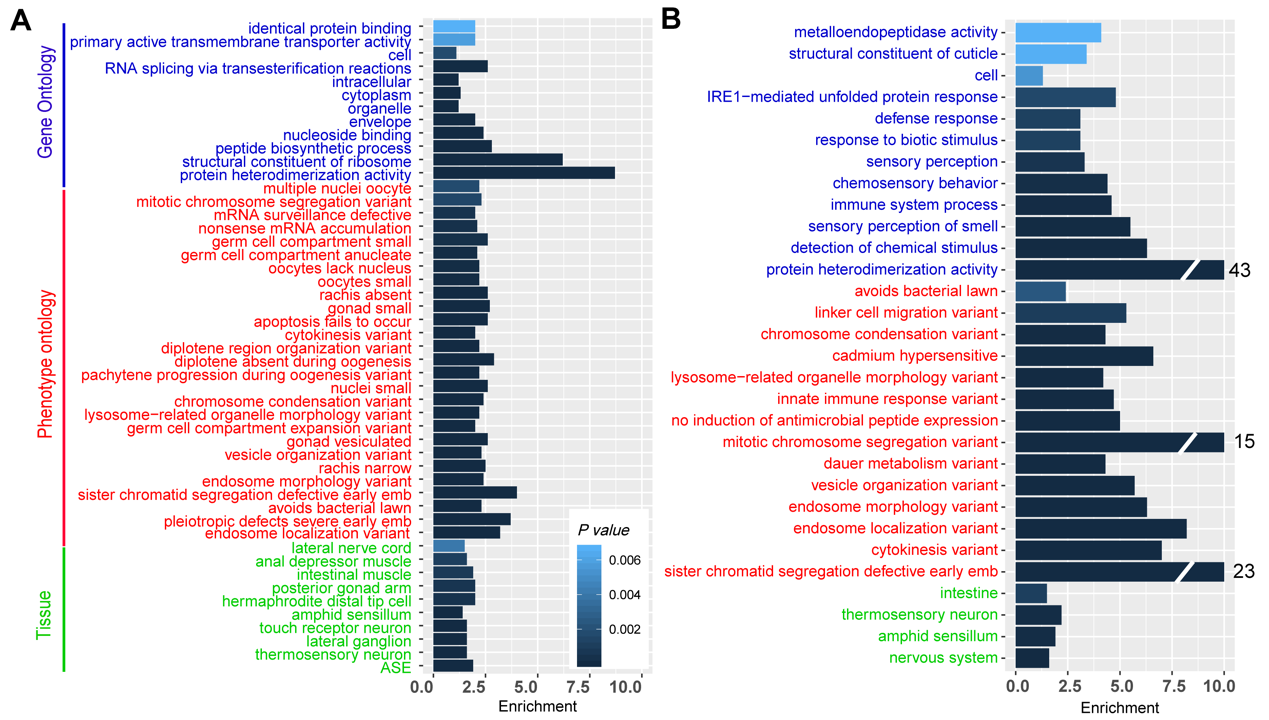

**Supplementary Figure S3.** Gene ontology enrichment analysis of genes with no nonsynonymous SNVs. (A) Gene enrichment analysis for the 1143 genes with no nonsynonymous SNVs and small indels in the CDS region. (B) Gene enrichment analysis for the 302 genes that have no nonsynonymous SNVs and small indels in the whole gene. Results of gene ontology enrichment (blue), phenotype enrichment (red), and tissue expression enrichment (green) were obtained using the enrichment analysis tools on Wormbase (WS275).

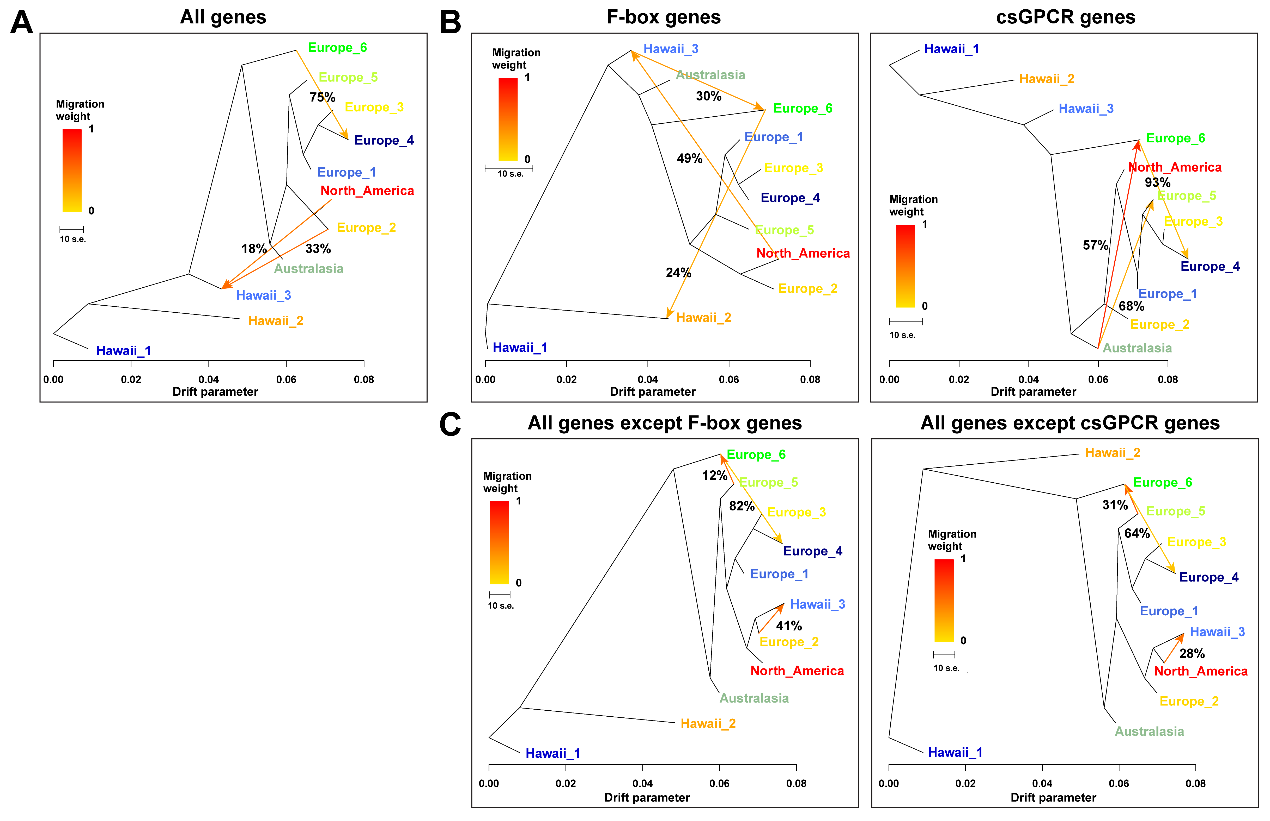

**Supplementary Figure S4. Gene flow of nonsynonymous variants of F-box and csGPCR genes among different subpopulations.** Gene flow among the three Hawaiian and eight non-Hawaiian subpopulations inferred by TreeMix using the nonsynonymous SNVs of all genes (A), the F-box genes, csGPCRs (B) and all genes excluding the F-box or csGPCR genes (C). Three migration events were allowed and 1000 bootstrap replicates were run with “Hawaii_1” set as the outgroup. The three most probable migration events from the 1000 bootstraps were shown and the percentage indicated the probability for the migration event. The color of the arrow indicates migration weight.

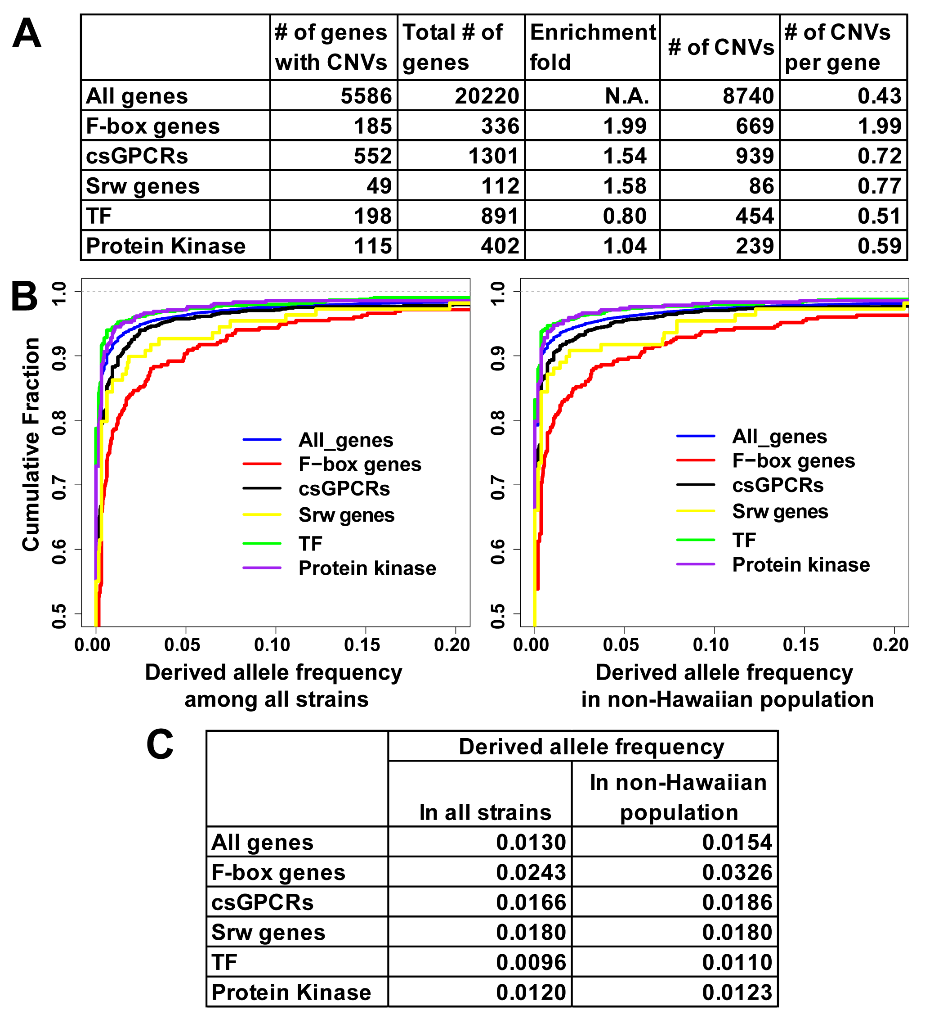

**Supplementary Figure S5. Polymorphism and derived allele frequency for copy number variants (CNVs) in the coding region of different families of genes.** (A) The number of genes with CNVs in coding region of all genes, F-box genes, csGPCRs, Srw genes, TF, and Protein kinase genes, as well as the number of CNVs occurring in these gene families. (B) The cumulative distribution of derived CNVs in different gene families in all strains or the non-Hawaiian population using XZ1516 as the outgroup. (C) The mean value of derived CNV allele frequency for different gene families.

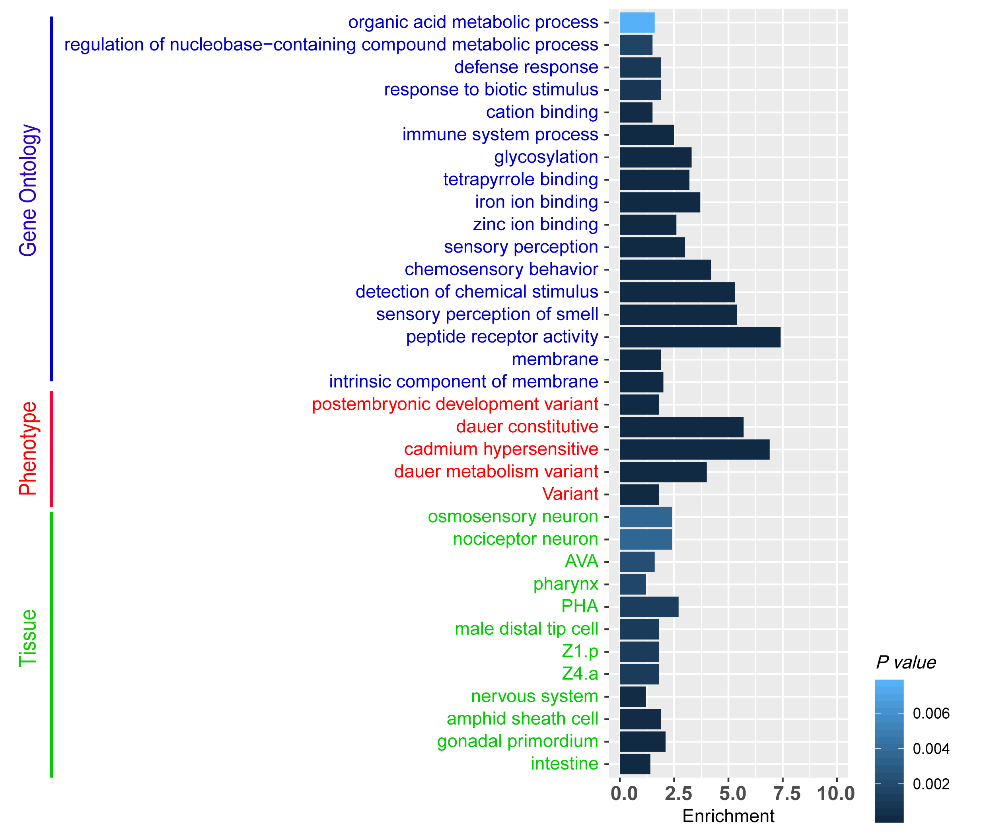

**Figure S6. Gene enrichment analysis for genes whose** **Fay and Wu’s *H* lower than -20.** Results of gene ontology enrichment (blue), phenotype enrichment (red), and tissue expression enrichment (green) were obtained using the enrichment analysis tools on WormBase (WS275).

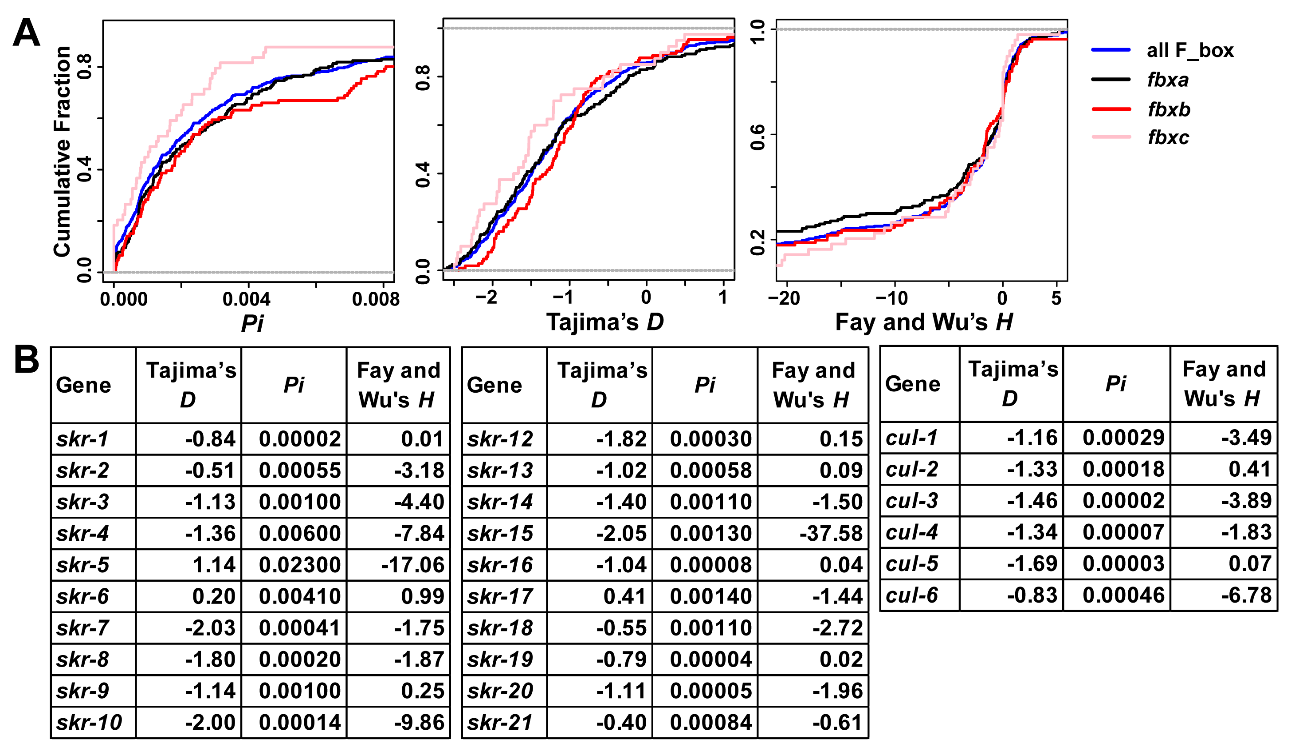

**Supplementary Figure S7.** **Population genetics analysis for SCF complex factors and genes in TF subgroups.** (A) The cumulative distribution of *Pi*, Tajima’s *D*, and Fay and Wu’s *H* values for *fbxa*, *fbxb*, and *fbxc* type F-box genes. (B) The *Pi*, Tajima’s *D*, and Fay and Wu’s *H* values of Skp1-homologus genes and Cullin genes. *skr-11* is a pseudogene and not shown. (C) The cumulative distribution of *Pi*, Tajima’s *D*, and Fay and Wu’s *H* value of genes in TF subgroups. See Table S1 for the classification of TF genes into subgroups. (D) The mean and median value of *Pi*, Tajima’s *D*, and Fay and Wu’s *H* values for all TFs and the five TF subgroups. Double asterisks indicate *p* < 0.05 in Wilcoxon’s rank-sum test.

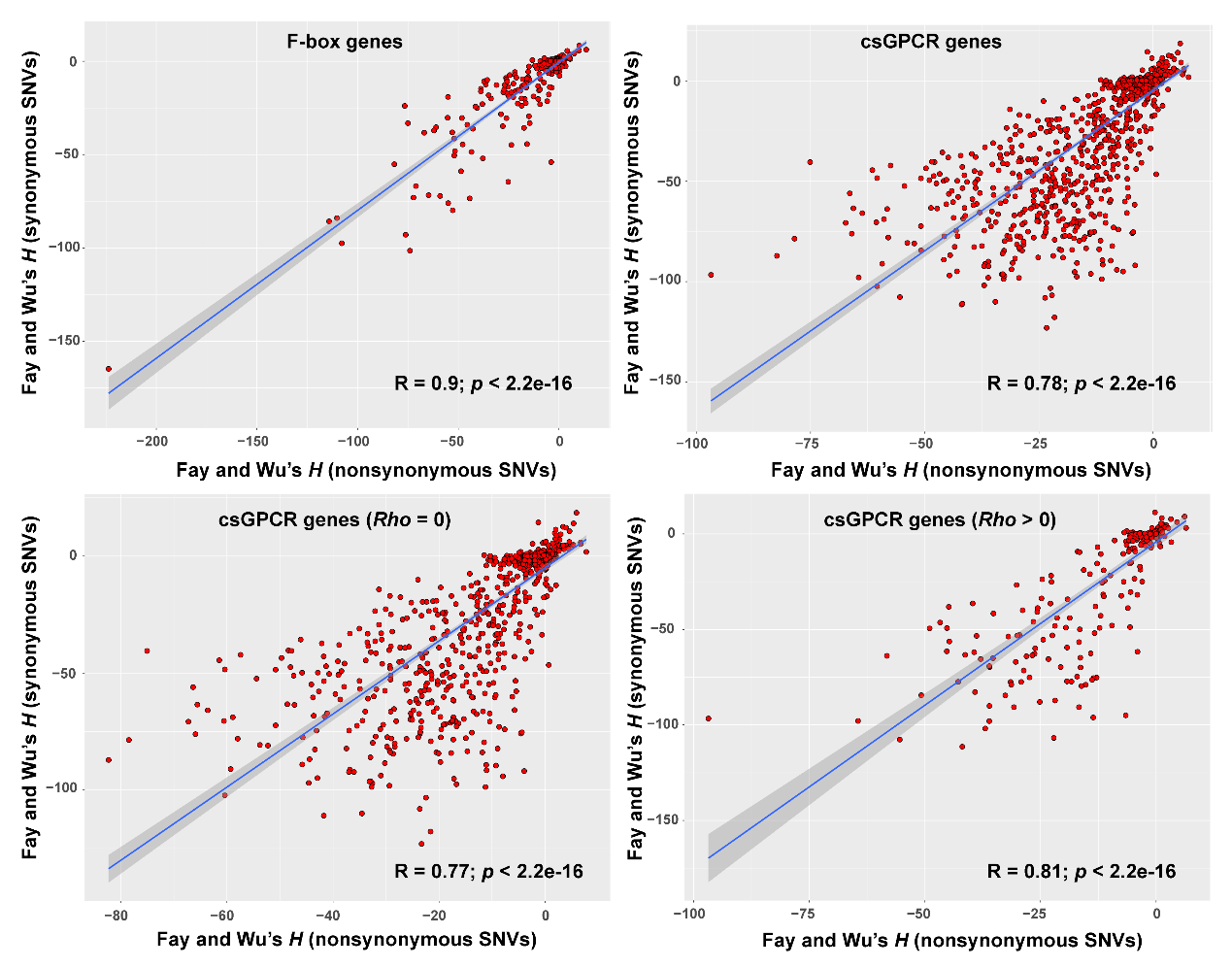

**Supplementary Figure S8. The correlation of Fay and Wu’s *H* values for nonsynonymous and synonymous SNVs of F-box and csGPCR genes.** The correlation of *H* values in low (*Rho* = 0) and high (*Rho* > 0) recombination regions is also shown for csGPCR genes.

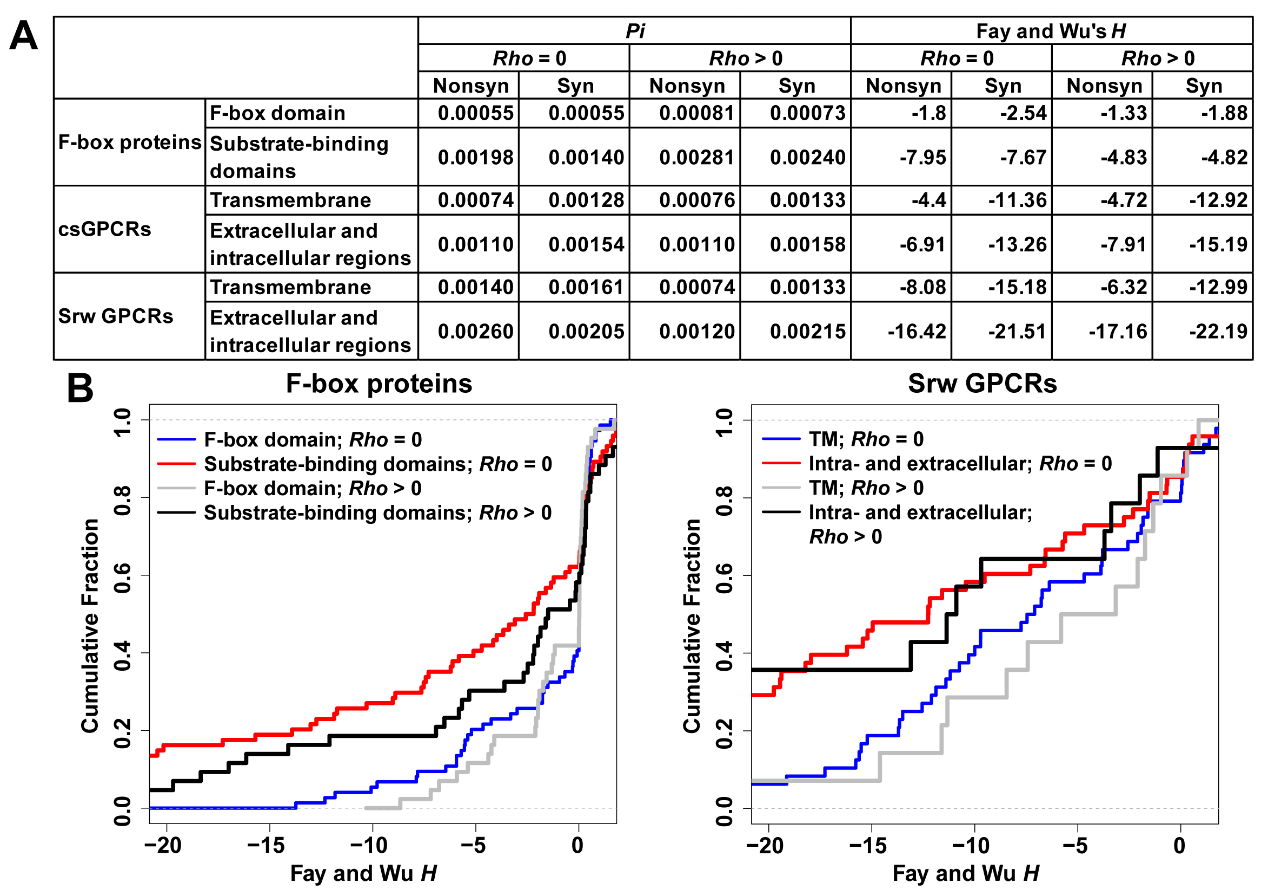

**Supplementary Figure S9. The *Pi* and Fay and Wu’s *H* values of nonsynonymous and synonymous SNVs in different domains of F-box, csGPCR and Srw proteins.** (A) The average of *Pi* and Fay and Wu’s *H* values for nonsynonymous and synonymous SNVs mapped to different domains of F-box, csGPCR and Srw proteins in low and high recombination regions. (B) The cumulative distribution of Fay and Wu’s *H* of nonsynonymous SNVs in F-box and substrate-binding domains of F-box proteins and the transmembrane (TM) and intra- and extracellular domains of Srw proteins in low and high recombination regions.

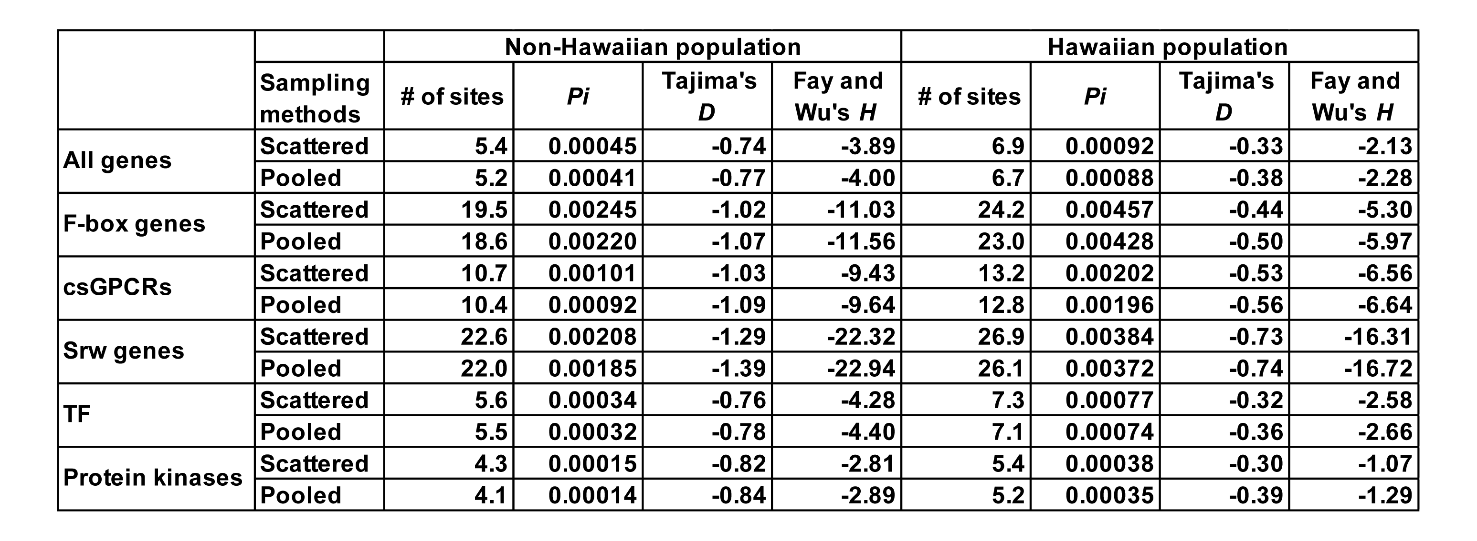

**Supplementary Figure S10. Polymorphisms and neutrality test statistics calculated using the nonsynonymous SNV data of strains randomly chosen under two different sampling methods.** Independent sampling were repeated 100 times to obtain 100 sets of strains and their SNVs. The mean of the number of segregating sites, *Pi*, *D*, and *H* for a given list of genes were calculated for each set of data. The average of these mean values are shown.

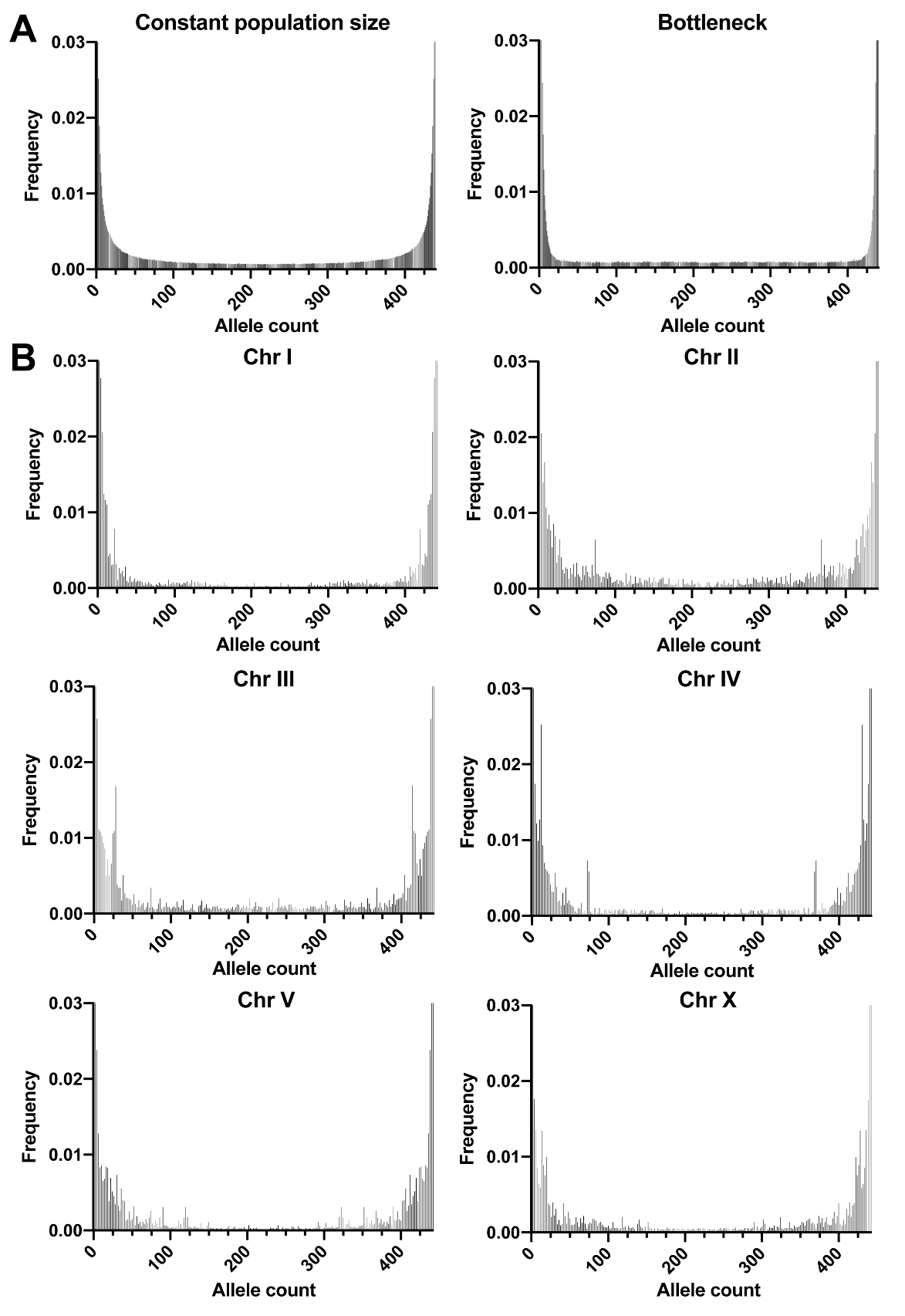

**Supplementary Figure S11. The site frequency spectrum (SFS) pattern.** (A) Site frequency spectrum of simulated SNV data under constant population size or bottleneck model. (B) The SFS pattern of the SNVs in the non-Hawaiian population from Chromosome I to X.

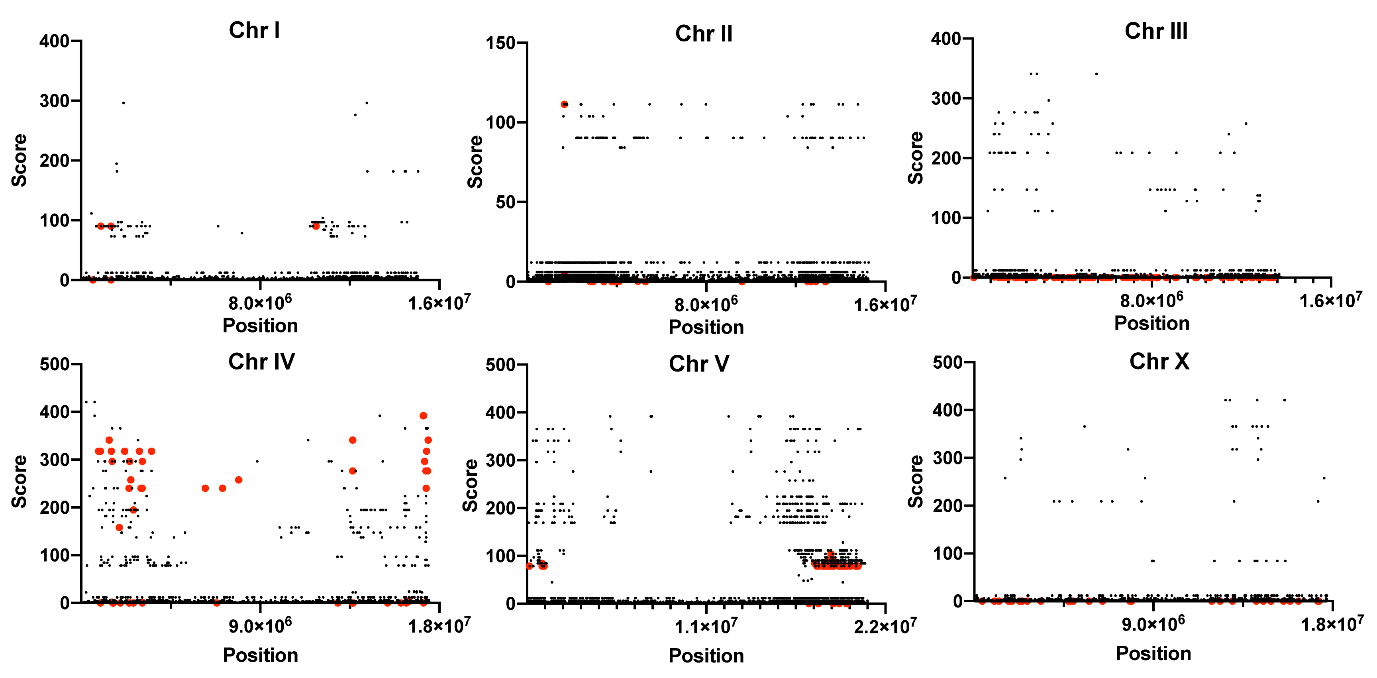

**Supplementary Figure S12. Selective sweep site prediction based on site frequency spectrum.** The distribution of α score predicted by SweeD for Chromosome I to X. X axis indicates chromosomal location. Red points indicate the alleles with significance level of 1%.

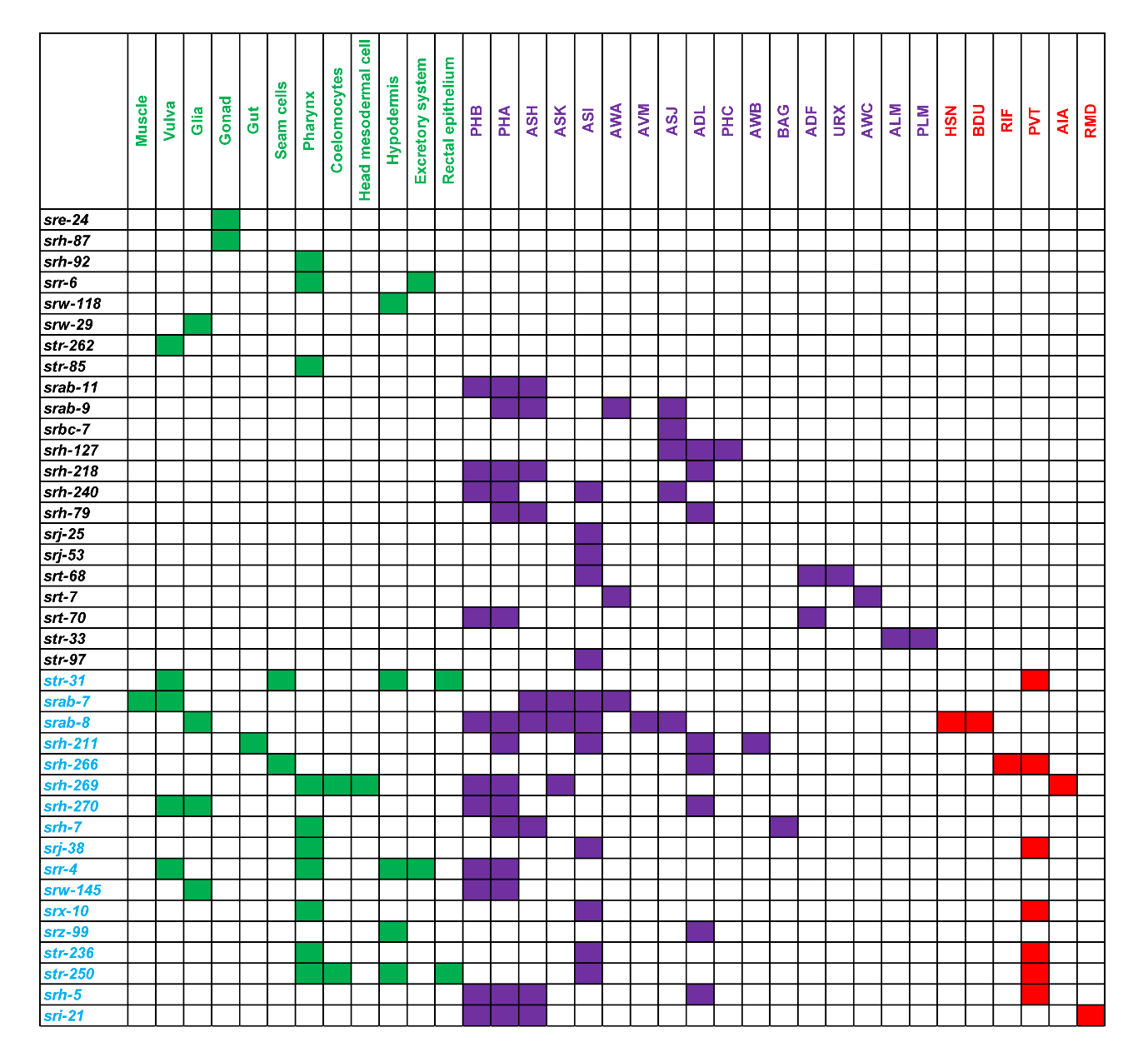

**Supplementary Figure S13. The expression atlas of some csGPCRs under strong positive selection.** The known expression pattern of csGPCR genes with Fay and Wu’s *H* value smaller than -20 according to Vidal, et al. (2018). Tissue types were divided into sensory (purple blocks), interneurons and motor neurons (red blocks) and non-neuronal tissues (green). Genes in black are expressed in one of the three types of tissues and genes in blue are expressed in more than one types of tissue.
